# Golgins support extracellular matrix secretion by collectively maintaining the Golgi structure-function relationship

**DOI:** 10.1101/2024.11.19.624265

**Authors:** George Thompson, Anna Hoyle, Philip A. Lewis, M. Esther Prada-Sanchez, Joe Swift, Kate Heesom, Martin Lowe, David Stephens, Nicola Stevenson

## Abstract

The secretion of extracellular matrix (ECM) proteins is vital to the maintenance of tissue health. One major control point of this process is the Golgi apparatus, whose dysfunction causes numerous connective tissue disorders. Golgi function is tightly linked to its structure, which is maintained by the cytoskeleton and Golgi organising proteins. We sought to investigate the role of two of these organising proteins, the golgins GMAP210 and Golgin-160, in ECM secretion. We found that loss of either protein had distinct impacts on Golgi organisation. GMAP210 loss caused cisternal fragmentation and dilation, alongside the accumulation of tubulovesicular structures. Meanwhile, Golgin-160 knockout lead to Golgi fragmentation and vesicle build-up. Nonetheless, loss of each protein had a similar impact on ECM secretion and glycosaminoglycan synthesis. We therefore propose that golgins are collectively required to create the correct physical-chemical space to support efficient ECM protein secretion and modification. This is the first time that Golgin-160 has been shown to be required for ECM secretion.

**Summary:** In this study, Thompson *et al* demonstrate that two cis-Golgi golgins, GMAP210 and Golgin-160, have distinct, non-redundant roles in maintaining Golgi organisation and that both are required to support the efficient secretion, assembly, and modification of extracellular matrix proteins.

## Introduction

The extracellular matrix (ECM) is a complex network of proteins, carbohydrate and mineral secreted by cells to build a scaffold that forms the structural basis of most tissues. To date, current resources describing the ‘matrisome’ have identified >1000 proteins associated with this compartment in humans (Shao et al., 2023), explaining the capacity for extensive diversification of biochemical and mechanical properties between tissue types. The most abundant family of ECM proteins are the collagens. These proteins assemble to form the main structural components of connective tissues alongside other fibre and network forming proteins such as fibronectin and elastin (Karamanos et al., 2021). This assembly is supported by accessory factors like the proteoglycans (Couchman and Pataki, 2012), a diverse set of proteins characterised by the presence of at least one glycosaminoglycan (GAG) chain which helps to promote tissue hydration and sequester signalling molecules (Gandhi and Mancera, 2008). Meanwhile ECM turnover is regulated by secreted proteases (Arpino et al., 2015; Lee and Murphy, 2004).

All newly synthesized ECM proteins must transit the secretory pathway to reach the extracellular environment for assembly. As they do so, many are heavily modified, which alters their chemistry and consequently the way in which they interact and assemble (Adams, 2023). Regulation of ECM protein modification is therefore crucial to the assembly of a functional matrix. One of the main compartments responsible for facilitating this process is the Golgi apparatus, which contains enzymes catalysing modifications such as O-glycosylation, N- and O-linked glycan maturation, proteolytic processing, phosphorylation, lipidation and sulphation (Potelle et al., 2015). Given this central role in protein processing it is unsurprising that a great number of genes affecting Golgi function have been linked to various connective tissue disorders (Hellicar et al., 2022).

Golgi structure is highly complex and tightly linked to its function. In mammalian systems it is composed of a stack of flattened, fenestrated membranous compartments called cisternae, which are laterally connected to form an interconnected Golgi ribbon (Lowe, 2011). These stacks are polarised such that cargoes arriving from the ER enter the Golgi stack at the *cis-*face, transit through the *medial* and *trans* compartments, and then get sorted before exiting the Golgi at the *trans* Golgi network (TGN). Golgi resident enzymes are distributed across the cisternae with a *cis-trans* polarity to support the successive addition of post-translational modifications to cargoes as they transit the compartments (Rabouille et al., 1995; Tie et al., 2022). This distribution is maintained by intra-Golgi vesicular transport which recycles enzymes between Golgi cisternae (Arab et al., 2024).

One of the major protein families responsible for maintaining Golgi organisation is the golgin family (Barr and Short, 2003; Gillingham and Munro, 2016; Witkos and Lowe, 2015). These proteins project coiled-coil domains into the cytosol from the Golgi surface to tether other Golgi membranes and transport vesicles (Lowe, 2019; Wong and Munro, 2014). Specific golgins localise to distinct regions of the Golgi organelle and this, combined with their affinity for different vesicles, confers directionality and specificity to intra-Golgi traffic (Wong et al., 2017). We and others have shown that one of these golgins, giantin, is required for the secretion of a healthy extracellular matrix. Loss of giantin prevents proper processing of the procollagen type I N-propeptide (Stevenson et al., 2021), impacts the abundance of ECM proteins (Katayama et al., 2018; Kikukawa and Suzuki, 1992), and impairs GAG metabolism (Kikukawa et al., 1991a; Kikukawa et al., 1991b; Kikukawa et al., 1990). These phenotypes are variable between animal models, but interestingly, always impact skeletal development (Katayama et al., 2011; Lan et al., 2016; Stevenson et al., 2017; Suzuki et al., 1988).

Another golgin which has been linked to ECM deposition and skeletal development is GMAP210. Mutations in the *TRIP11* gene encoding GMAP210 are causative of the human skeletal diseases achondrogenesis type 1A and odontochondrodysplasia (Costantini et al., 2021; Del Pino et al., 2021; Medina et al., 2020; Qian et al., 2021; Smits et al., 2010; Upadhyai et al., 2021; Wehrle et al., 2019; Yeter et al., 2022). Interestingly, *Trip11* mutant mice have a similar phenotype to giantin mutant rats, with both models showing neonatal lethal chondrodysplasia characterised by craniofacial defects, shortened limbs and ribs, and delayed mineralisation of bone (Bird et al., 2018; Follit et al., 2008; Smits et al., 2010; Yamaguchi et al., 2021). Loss of GMAP210 also impacts the secretion of specific ECM proteins in a selective manner (Bird et al., 2018; Smits et al., 2010; Yamaguchi et al., 2021)

Altogether these studies suggest that extracellular matrix secretion is particularly susceptible to the loss of at least two golgins, which appear to act non-redundantly in this context. This led us to question the extent to which ECM deposition is sensitive to the loss of golgins more widely, and how this relates to changes in Golgi organisation. To begin addressing this, we performed a comparative study between GMAP210 and a third golgin, Golgin-160. Golgin-160 was chosen because it resides on *cis-*Golgi membranes like GMAP210 (Hicks et al., 2006) but remains poorly characterised at a functional level. Here we report that knockout of Golgin-160 has a similar impact on ECM organisation and composition as GMAP210 loss. Golgi morphology on the other hand, is affected differently, consistent with the two golgins acting in different ways to maintain Golgi homeostasis. We therefore conclude that GMAP210 and Golgin-160 function non-redundantly to create the correct physical and biochemical space to support the modification and secretion of complex ECM molecules.

## Results

### Generation of golgin knockout cell lines

To test the role of different golgins on matrix secretion, we generated two GMAP210 and Golgin-160 knockout (KO) clones using CRISPR-Cas9 genome engineering. GMAP210 KO clone 1 was generated using a gRNA targeting exon 4 of the *TRIP11* gene (Supplemental Figure 1A). This exon is common to both predicted protein coding transcripts TRIP11-201 (Ensembl transcript ID <u>ENST00000267622.8</u> Uniprot ID <u>Q15643-1</u>) and TRIP11-202 (<u>ENST00000554357.5</u>, <u>H0YJ97</u>) and so mutation should impact all protein expression. The generated mutation was a single base pair insertion of alanine resulting in a frame shift from position Glu^114^ and truncation at amino acid 155 (Supplemental Figure 1A). GMAP210 KO clone 2 was generated using a gRNA targeting exon 1 of transcript TRIP11-202 and upstream of the predicted start site of the longer transcript TRIP11-201. This resulted in a 10bp deletion in the promoter region (Supplemental Figure 1A). Loss of protein expression was confirmed in both clones by immunofluorescence using three different antibodies targeting the N- and C-terminus of the GMAP210 protein (Supplemental Figure 1B) and western blotting (Supplemental Figure 1C). Note trance amounts of protein could be detected with the C-terminal antibody suggesting a truncation product may persist. Nonetheless, we considered this sufficient depletion for further study.

Golgin-160 KO clones were generated using a gRNA targeting exon 20 of transcript GOLGA3-201 (ensemble ID <u>ENST00000204726.9</u>, Uniprot ID: <u>Q08378-1</u>). This is expected to mutate all predicted transcripts encoding fragments larger than 44 amino acids. Mutation with this gRNA consistently produced a single alanine insertion, resulting in frame shift at amino acid Glu^1312^ and truncation at amino acid 1331 (Supplemental Figure 1A). Consequently, two clones with the same mutation were carried forward for experiments to rule out clonal effects. Protein loss was confirmed by immunofluorescence and western blotting, again with antibodies targeting both the N- and C-terminus (Supplemental Figure 1D,E).

### Loss of either GMAP210 or Golgin-160 causes changes in ECM assembly and organisation

To investigate the requirement for GMAP210 and Golgin-160 in ECM secretion and assembly, we grew our new KO lines for a week in the presence of ascorbic acid to promote collagen secretion and visualised the cell-derived ECM by immuno-fluorescence labelling. Co-labelling of two major fibrillar ECM components, collagen type 1 and fibronectin, revealed an extensive network of highly aligned and organised extracellular fibrils in wildtype (WT) cell cultures, with cell nuclei stretching along the fibril axis (Figure 1A-D). In GMAP210 (Figure 1A-B) and Golgin-160 KO (Figure 1C-D) cell cultures on the other hand, fibrils were identifiable, but they were less well ordered, showing poor alignment and an increase in branching. To examine the structure of these fibrils more closely, the cell layer was extracted, and the remaining cell-derived matrix was imaged by high-speed atomic force microscopy (HS-AFM). Again, fibrils were less well organised in all golgin KO cultures (Figure 1E-F). They were also thinner, however the periodicity of the collagen fibrils was largely unchanged, suggesting assembly of collagen trimers into fibrils is normal but lateral interconnections between fibrils may be reduced.

**Figure 1.**
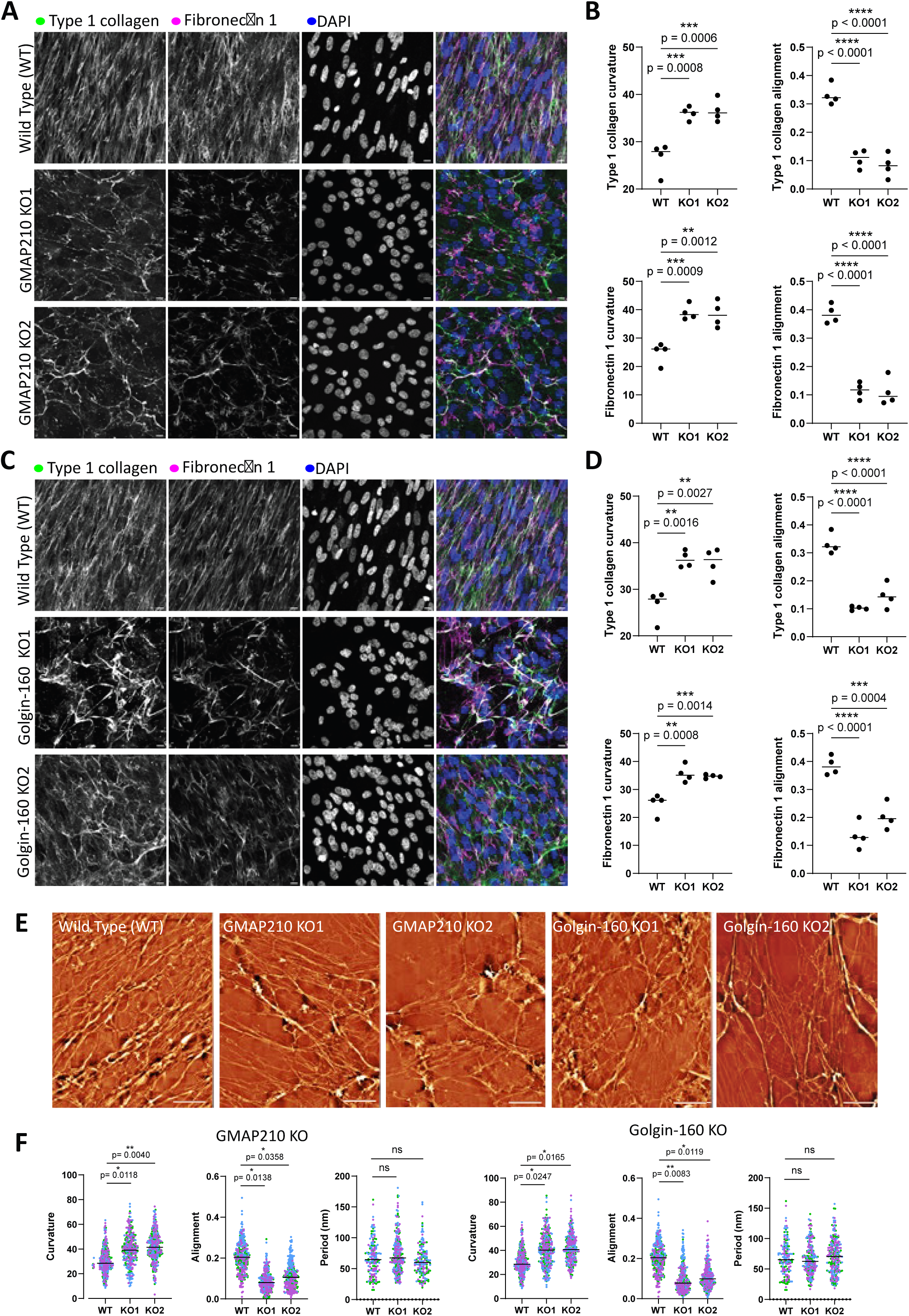
Extracellular matrix organisation is altered in golgin mutant cultures. **A, C.** Confocal maximum projection images of unpermeabilised WT, GMAP210 KO (A) and Golgin-160 KO (C) RPE1 cell cultures immunolabelled for extracellular collagen type I (green), fibronectin-1 (magenta) and nuclei (DAPI, blue). Scale bars 10 µm. **B, D.** Quantification of fibril characteristics measured from images represented in (A,C). Individual dots represent mean of each biological replicate (n=4) and bars represent median of all experiments. **E.** Decellularised ECM from WT, GMAP210 KO and Golgin-160 KO cell cultures imaged by HS-AFM. Images are representative 10 x 10 raster scans from biological replicates (n=3). Scale bars 10 µm. **F.** Quantification of fibril characteristics measured from images represented in (E). Individual dots are measurements from each tile in the raster scan, with each biological replicate colour coded (n=4). Bars represent median of all experiments. **B, D, F.** Data were subjected to a Shapiro Wilk test for normality and then a nested one way analysis of variance (ANOVA) with Dunnet’s test for multiple comparisons to generate p values.

The overall fluorescence intensity of collagen staining in the matrix appeared reduced in the KO cultures. We therefore determined the abundance of collagen type I in lysate and matrix fractions of cell cultures by immunoblotting against the Col1a1 chain. Col1a1 abundance in the lysate fractions of mutant cultures was comparable to that of WT lysates when normalised to total protein, suggesting expression is normal and collagen is not retained inside the cell (Figure 2A,Ci). Col1a1 abundance was, however, reduced in the ECM fraction of all mutant lines when normalised against total cellular protein (Figure 2B, Cii), suggesting it is not being efficiently incorporated into the insoluble matrix. Collagen in the media was below the level of detection by this method.

**Figure 2.**
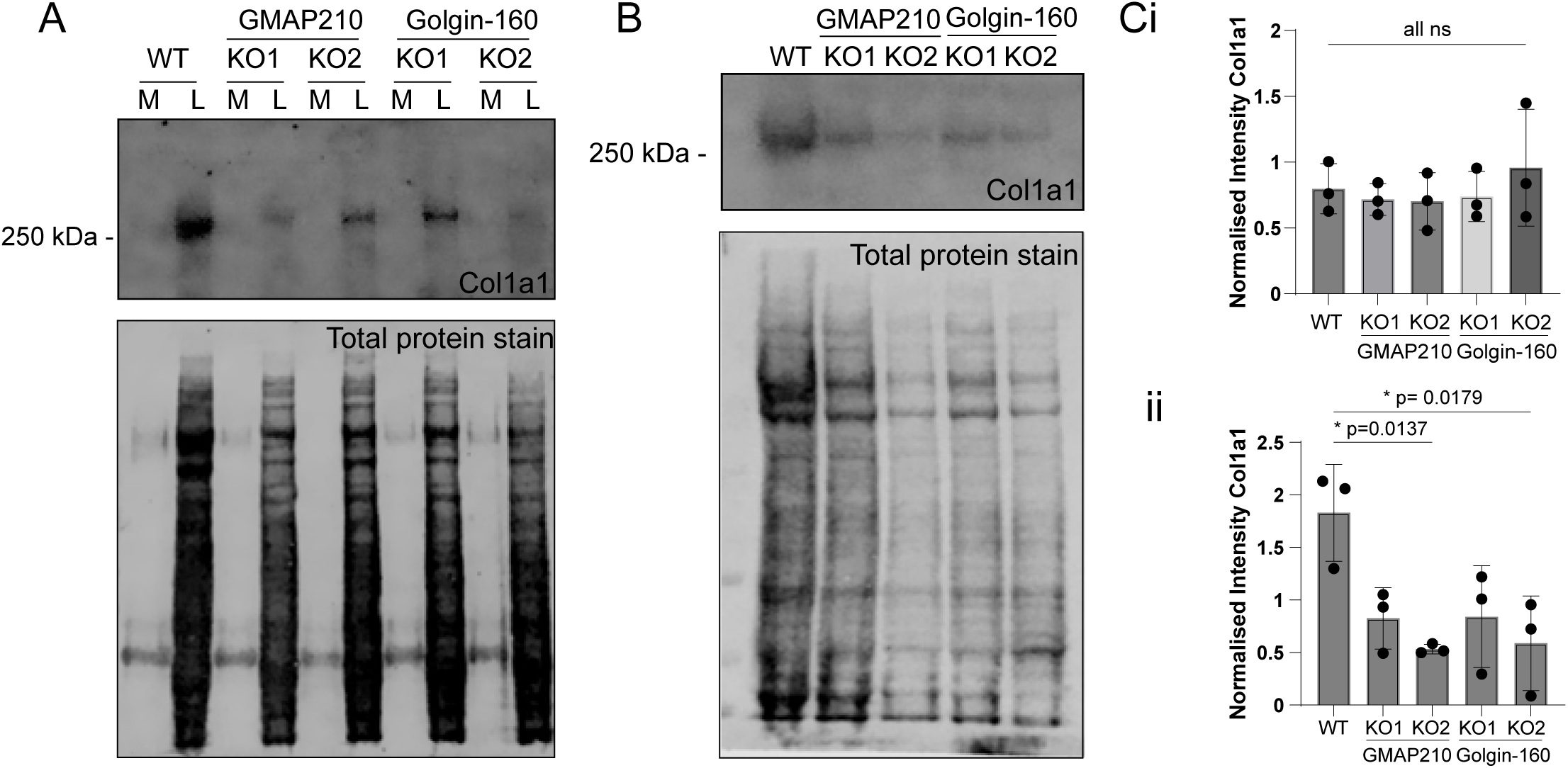
Collagen deposition in the matrix is impaired in golgin mutant cells. **A.** Immunoblot for Col1a1 and total protein stain after SDS-PAGE of medium (M) and lysate (L) samples taken from wildtype and golgin mutant cultures. **B.** Immunoblot for Col1a1 and total protein stain after SDS-PAGE of the cell derived matrix extracted from wildtype and golgin mutant cultures. **C.** Quantification of Col1a1 intensity normalised against total cellular protein for **i.** lysates samples as represented in A and **ii.** matrix samples as represented in B. Dots show individual experiment result (n=3), bars show mean and standard deviation. Data were subjected to a Shapiro Wilk test for normality and then a nested one-way ANOVA with Dunnet’s test for multiple comparisons to generate p values.

### ECM composition is altered following loss of GMAP210 and Golgin-160

To determine whether the observed changes in ECM organisation in the golgin KO cultures reflect changes in ECM composition, the cell-derived matrix from WT and KO cultures was collected and analysed by mass-spectrometry. Principal component analysis showed distinction between the WT and Golgin-160 KO and between the KO matrix samples, but variation between repeats was high (Supplementary Figure2A). Results were filtered against the matrisome database (Shao et al., 2023) to indicate proteins specifically related to ECM structure and function. Overall, fewer proteins were significantly impacted by loss of Golgin-160 than of GMAP210, however there was a great deal of overlap between the KO lines (Figure 3). For example, laminin subunit gamma 1 (LAMC1), Laminin subunit alpha 5 (LAMA5), hemicentin-1 (HMCN1), Latent Transforming Growth Factor Beta Binding Protein 1 (LTBP1) and Transforming Growth Factor Beta 2 (TGFB2) were significantly reduced in abundance in at least one Golgin-160 KO line as well as the GMAP210 KOs (Figure 3A-C). Conversely, the abundance of fibrillins was increased in all KO lines (Figure 3A-C). Interestingly, Lysyl Oxidase Like 1 (LOXL1), which was upregulated in the GMAP210 KO ECM, was downregulated in Golgin-160 KOs, and vice versa for Procollagen-Lysine,2-Oxoglutarate 5-Dioxygenase 1 (PLOD1, Figure 3A-C). This hints at golgin specific changes in the post-translational modification of collagen. Altogether these data show that both GMAP210 and Golgin-160 are required for the deposition of a well-organised ECM. The similarity seen between ECM phenotypes in GMAP210 and Golgin-160 KO cells also demonstrates the non-redundant nature of these golgins in this context and the broad susceptibility of ECM cargoes to golgin loss.

**Figure 3.**
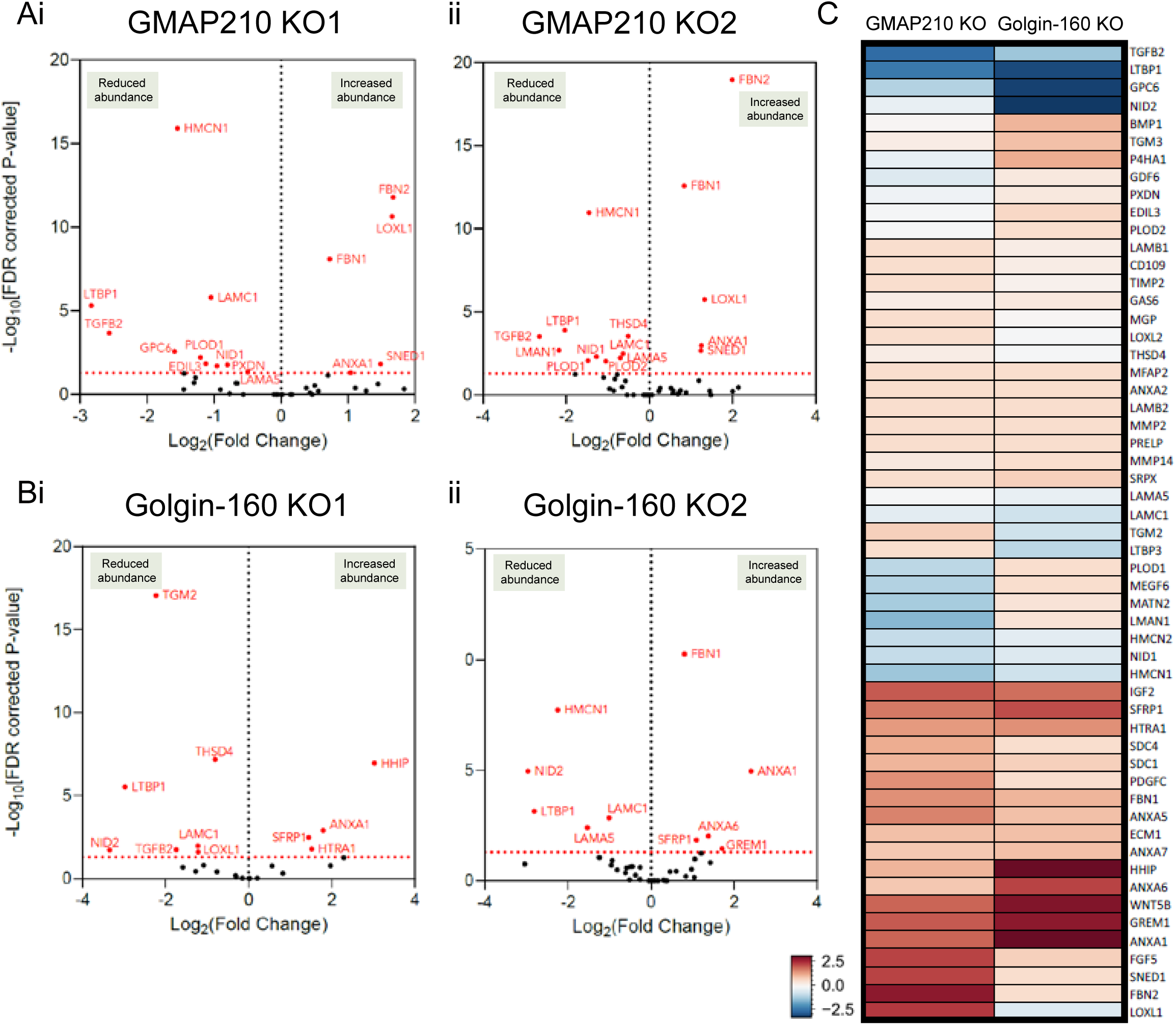
Extracellular matrix composition is altered in golgin KO cells. **A-B.** Volcano plots showing log-fold change in cell-derived matrix protein abundance from GMAP210 KO (A) or Golgin-160 KO (B) cells compared to WT. Red-labelled points represent proteins with significantly changing abundance, with Benjamini-Hochberg false discover rate (FDR) corrected *p*- values < 0.05, n=3. Only proteins categorised as ‘matrisome’ are represented on these plots (Naba 2011). Results for two different clones per gene are shown (i,ii). **C.** Heat map comparing protein abundance changes between the two different golgin KOs after pooling both clones. Red and blue indicate increased and decreased abundance respectively. Colour intensity is determined by average log-fold change across mutants compared to WT.

### GMAP210 and Golgin-160 are not required for procollagen processing

We have shown previously that KO of giantin disrupts procollagen processing (Stevenson et al., 2021). To test whether impaired collagen processing is contributing to the ECM phenotypes reported here too, we interrogated our ECM proteomics data set for peptides relating to collagen pro-domains. Collagen type IV had the highest peptide count and therefore provided the most robust analysis. Comparing the average peptide counts across the Col4a2 sequence we found no significant increase in the number of peptides containing an intact N-propeptide cleavage site in the KO cultures (N-propeptide overlap sequence, Supplemental Figure 2B and C), indicating that processing of collagen type IV proceeds effectively in the absence of Golgin-160 and GMAP210. Expression of a procollagen type 1 construct in which the N-propeptide is GFP tagged (McCaughey et al., 2019), also revealed the presence of free collagen type I N-propeptide cleavage products in both WT and Golgin-160 KO cells indicating successful cleavage (Supplemental Figure 2D). GMAP210 KO cells did not tolerate procollagen type I over-expression and could not be tested. Altogether, these data indicate that loss of GMAP210 or Golgin-160 does not phenocopy loss of giantin with respect to collagen processing, and that this is not a contributing factor to the observed ECM phenotypes here.

### Loss of GMAP210 or Golgin-160 leads to distinct changes in Golgi morphology

Having established that loss of GMAP210 or Golgin-160 has a similar impact on ECM deposition, we next sought to determine the impact of golgin loss on Golgi organisation. Cells were labelled with markers of the *cis*- and *medial*-Golgi and the *trans*-Golgi network (TGN). Golgi area and extent of fragmentation were then measured (Figure 4A-D). Strikingly, each golgin KO had a different impact on Golgi morphology. As previously reported (Bird et al., 2018; Sato et al., 2015; Smits et al., 2010; Wehrle et al., 2019; Yamaguchi et al., 2022), loss of GMAP210 resulted in a clear compaction of Golgi structures compared to wildtype cells (Figure 4A-B). The Golgi also appeared fragmented, although this was difficult to quantify due to this compaction and poor definition of Golgi elements. In Golgin-160 KO cells on the other hand, no consistent change in Golgi size was observed, but there was a clear increase in Golgi fragment number (Figure 4C-D). In all KO lines, the *cis-trans* polarity of the Golgi appeared to be maintained, although again this was harder to ascertain in the GMAP210 KO cells (Figure 4A, C).

**Figure 4.**
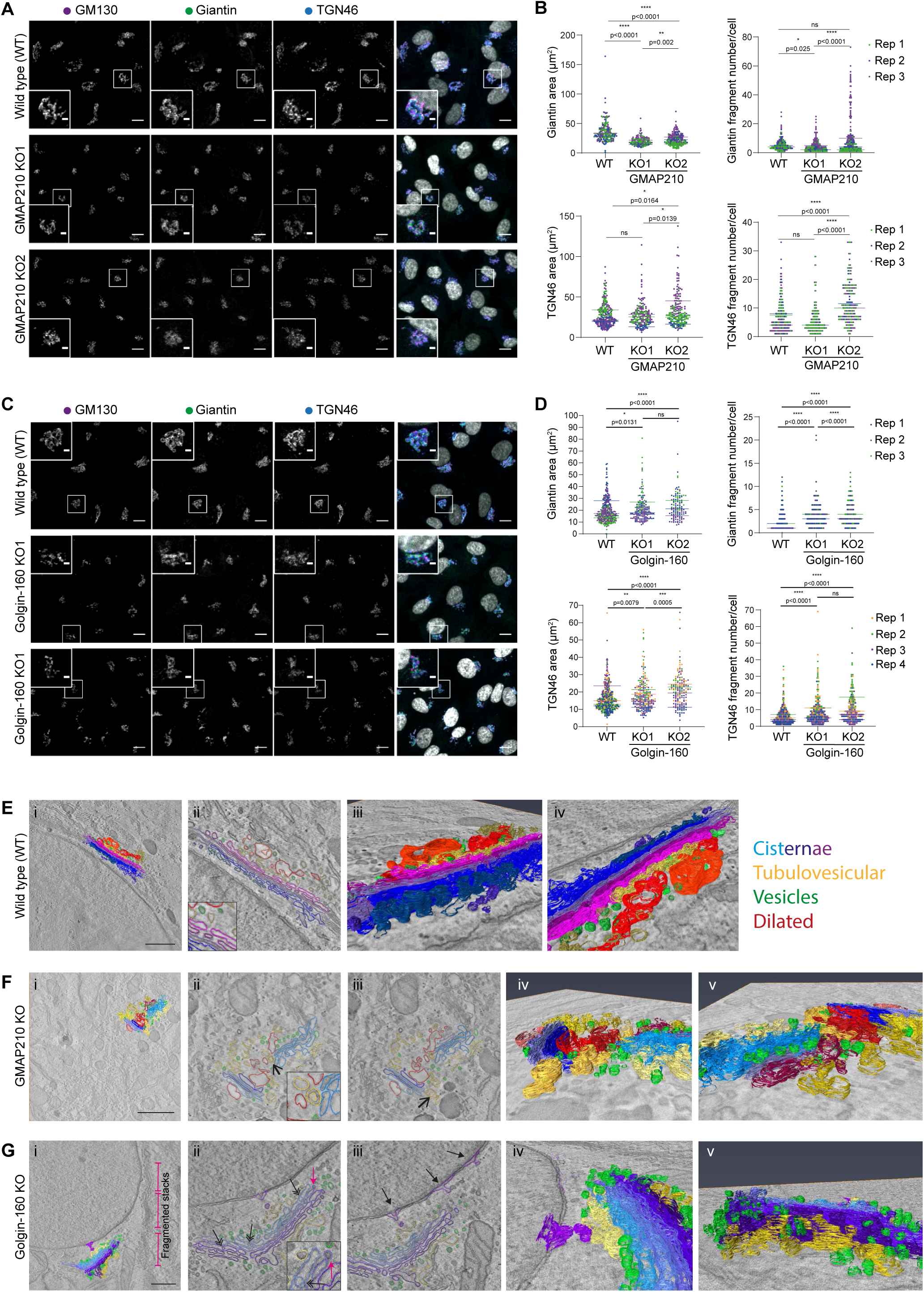
Golgi organisation is altered upon loss of GMAP210 or Golgin-160. **A, C.** Maximum projection confocal images of WT and GMAP210 KO (A) and Golgin-160 KO (C) RPE1 cells immunolabelled for cis-Golgi (GM130, magenta), cis-medial-Golgi (Giantin, green) and trans Golgi network (TGN46, blue) markers. Nuclei labelled with DAPI (greyscale). Scale bar 10 µm. Inset scale bar 1 µm. **B, D.** Quantification of total giantin and TGN46 area and fragment number per cell from images represented in (A,C). Individual dots represent one cell and are coloured by replicate (n=3). Bars show the median from each replicate experiment. Statistical analysis performed using Shapiro-Wilk normality test and Kruskall-Wallis significance test. **E-G.** Tomographic reconstructions of Golgi structures in WT (E), GMAP210 KO (F), and Golgin-160 KO (G) cells. Segmented membranes are labelled as cisternae (blue/purple), dilated structures (red), tubulovesicular structures (yellow), and vesicles (green). **Eii, Fii-iii, Gii-iii** Single slice images with segmentation. **Fii-iii** Open arrows point to invaginations within spherical regions of tubulovesicular structures. **Gii-iii** Double headed arrows indicate top to bottom fenestrations in cisternae, closed headed arrows indicate budding structures at the nuclear envelope, pink arrow shows frustrated budding/fusion intermediate. **Ei, iii-iv, Fi, iv, v, Gi, iv, v.** 3D rendering of segmentation. **E-Gi.** Scale bar 1 µm.

To better define the impact of KO on Golgi membrane organisation, higher resolution imaging was performed using single-tilt electron tomography. For each cell line, 3D tomographic reconstructions were generated (Supplemental movies 1-3) and a section of the Golgi was segmented to build a 3D model of membrane organisation (Supplemental movies 1-3, Figure 4E-G). In these models, membranes associated with the Golgi apparatus were grouped into four structural categories: cisternae (blue), defined as laterally flattened, stacked membranous compartments; dilated structures (red) - large volume structures lacking flattened regions; tubulovesicular structures (yellow) - convoluted structures with interconnected regions of spherical and tubular shape; and vesicles (green) – small, spherical structures.

In wildtype cells, as expected, we observed tightly stacked, evenly flattened, fenestrated cisternae that were laterally connected to span lengths of tens of microns (Figure 4E). Dilated and tubulovesicular structures were predominantly localised to the *trans* side of the Golgi in these cells, with small numbers of vesicles concentrated at the cisternal rims and *trans*-Golgi face. In contrast, although the Golgi was identifiable in GMAP210 KO cells, membranes were highly disordered (Figure 4F). The cisternae were mostly stacked, but they were short with no lateral connections and few fenestrations, and many were dilated or had bulges along their length, particularly at the *cis*- and *trans*- end of the stack. Some membranous structures resembling these dilated cisternae were also observed near, but not part of, a stack but their origin and identity is hard to assign. Finally, in many instances the cisternae were surrounded by extensive tubulovesicular structures interspersed with vesicles that wrapped around the mini-stacks (Figure 4Fiii, iv). Some of these structures had tunnel like indentations, which in cross-section appeared as internalised vesicles but were continuous with the cytosol (Figure 4Fii-iii, open headed arrows). Note, we did not see any obvious ER swelling as has been reported to occur in a cell type specific manner in some *in vivo* studies (Bird et al., 2018; Smits et al., 2010; Yamaguchi et al., 2021). Despite the similarity of the tubulovesicular membranes to ERGIC membranes (Appenzeller-Herzog and Hauri, 2006), we did not see any gross changes to ERGIC53 distribution in these cells at the light level (Supplementary Figure 3A). To conclude, the geometry and organisation of Golgi membrane is severely perturbed in the absence of GMAP210 in a way that is suggestive of both structural and trafficking defects.

Examination of Golgin-160 KO cells showed well stacked, tightly flattened cisternae that remained juxtanuclear as in wildtype cells (Figure 4G). However, consistent with the immunofluorescence data, there were very few lateral connections between cisternae meaning membranes were fragmented into closely apposed ministacks rather than forming a classical ribbon architecture (Figure 4Gi, pink brackets). Cisternae also appeared to have internal vesicles or holes unlike control cells (Figure 4Gii, double headed arrows). Strikingly, there was a clear accumulation of vesicular structures in the area surrounding the cisternal rims, as well as both the *cis*- and *trans*-Golgi face (Figure 4Gv). This coincided with a high incidence of spherical structures attached to the membranes of cisternal rims that resembled frustrated vesicle budding or fusion profiles (Figure 4Gii, pink arrow). Intriguingly, budding vesicles were also frequently observed at the nuclear envelope (Figure 4Giii, closed head arrows). Although the accumulating vesicles were reminiscent of COPI vesicles with an electron dense coat, we were unable to observe any gross changes in COPI localisation at the light level by immunofluorescence and individual puncta could not be distinguished in the Golgi region (Supplemental Figure 3B). In conclusion, loss of GMAP210 or Golgin-160 has a different impact on Golgi structural organisation.

### Loss of either GMAP210 or Golgin-160 leads to reduced secretion of ECM components and regulators

Given the clear disruption to Golgi organisation in our knockout lines, we decided to look at secretion more generally. Cells were grown in serum-free medium overnight, before collecting the media to analyse the abundance of soluble secreted proteins by mass spectrometry. Three independent replicate samples were collected for all WT and KO lines and analysed simultaneously by 15-plex tandem-mass tagging mass spectrometry. The sum of raw abundance revealed replicate one to have contained more protein and thus pair-wise comparisons within replicates were chosen over averages (Supplemental Figure 4A). Principal component analysis reiterated the need for paired analysis with PC1 and 2 separating the samples by replicate, however good separation was observed at PC3 and PC4 between WT, Golgin-160 KO and GMAP210 KO lines, with individual clones clustering together for each KO. This indicates genotype is a key contributor to variance and highlights variation between secretomes in each line (Supplemental Figure 4B). The two individual KO clones for each golgin were pooled using a linear regression model to increase statistical power. Log-fold change in protein abundance was plotted against p-value to identify proteins whose secretion is significantly altered in KO cultures compared to WT (Figure 5).

**Figure 5.**
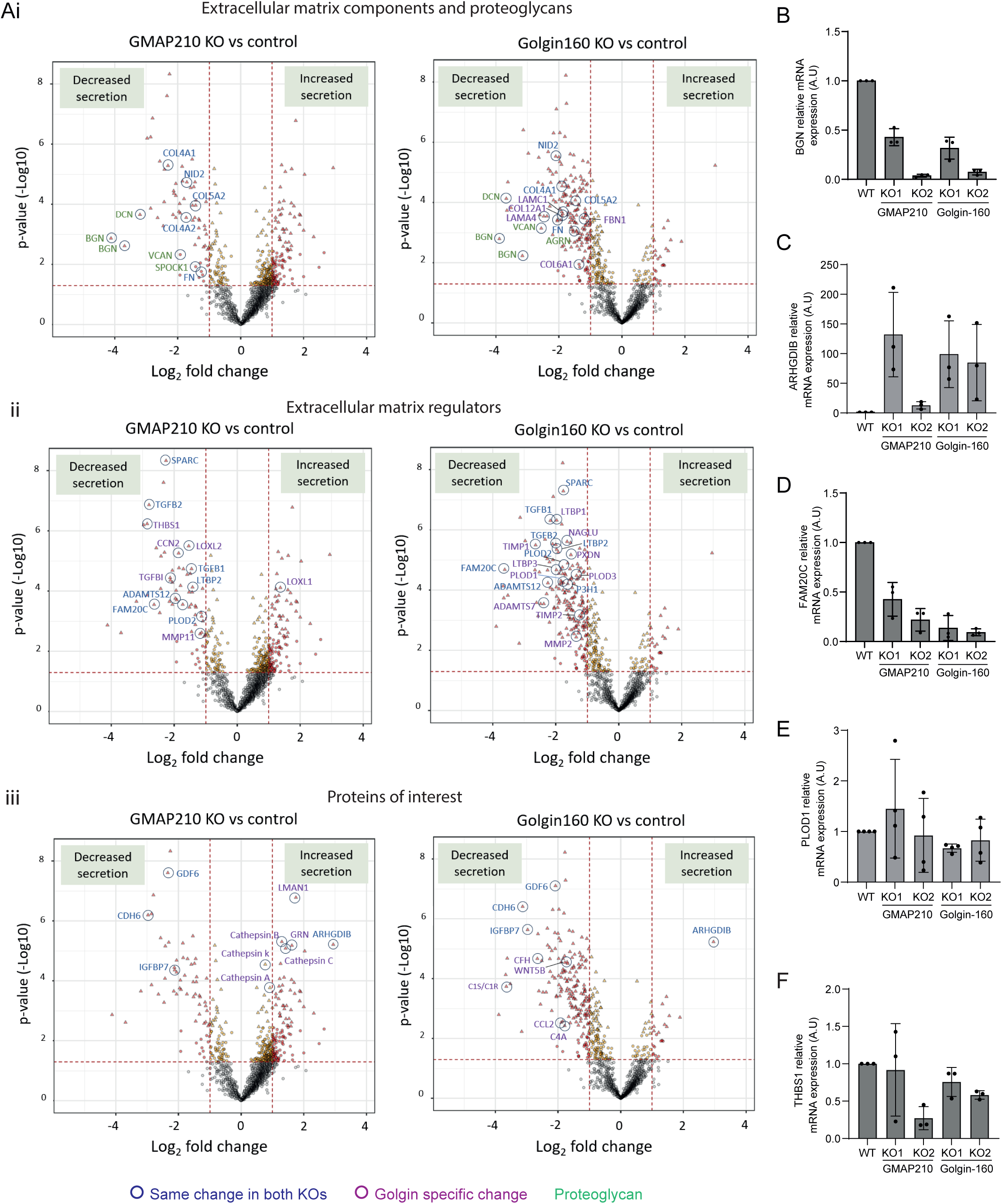
General secretion is differentially impacted by loss of GMAP210 or Golgin-160. **A.** Volcano plots showing log-fold change in protein abundance in the media from KO versus WT cultures plotted against significance across three independent replicate experiments. The results from the two golgin KO clonal cell lines were pooled to increase power. Lines and dot colours on the graph are for p<0.06 and >1 or <-1 LogFC. Triangular points show hits which pass an FDR corrected p<0.05. Structural ECM proteins (i), ECM regulators (ii) and other proteins of interest (iii) have been highlighted. Proteins significantly changed in the same way in both KO cultures are labelled in dark blue whilst proteins impacted in a golgin specific way are labelled in purple. The proteoglycan family is indicated by green text. **B-F.** RT-PCR analysis of gene expression of (B) BGN, (C) ARHGDIB, (D) FAM20C, (E) PLOD1, and (F) THBS1. Expression levels normalised to expression of housekeeping gene YWHAZ. Individual data points represent biological replicate experiments (n=3-4). Bars show mean and standard deviation.

In total, 482 proteins and 427 proteins showed altered raw abundance (p<0.05) in the secretome of GMAP210 KO and Golgin-160 KO cultures respectively compared to wild-type cells (Figure 5A, Supplemental Figure 4C). GO term analysis revealed that the top 15 biological pathways affected in both golgin KO samples included collagen fibril and extracellular matrix organisation, consistent with the ECM defects described above, as well as cell adhesion and migration (Supplemental Table 1). Proteins involved in osteoblast and chondrocyte differentiation were also impacted by golgin loss. This was not surprising for the GMAP210 KOs given the skeletal phenotypes reported in animal models and human disease (Bird et al., 2018; Costantini et al., 2021; Del Pino et al., 2021; Follit et al., 2008; Medina et al., 2020; Qian et al., 2021; Smits et al., 2010; Upadhyai et al., 2021; Wehrle et al., 2019; Yamaguchi et al., 2021), but intriguingly, a similar effect was seen following loss of Golgin-160, which has not been reported to cause a skeletal phenotype in mice (Banu et al., 2002; Bentson et al., 2013). Pathways associated with proteolysis and signalling appeared to be impacted by the loss of Golgin-160 specifically (Supplemental Table 1).

Examination of the individual proteins most heavily affected by loss of each golgin shows a clear depletion of ECM components and regulators in the media of both KO cultures, with a high degree of overlap between GMAP210 and Golgin-160 KOs (Figure 5A - proteins impacted in the same way by each KO circled blue, Supplemental Figure 4C). For example, the protein family showing the greatest depletion in both mutant media fractions were the proteoglycans. Biglycan (BGN), decorin (DCN), versican (VCAN) all showed reduced abundance in both KO samples, with aggrecan (AGRN) also depleted in the Golgin-160 KO and testican (SPOCK1) impacted in the GMAP210 KOs. This suggests the proteoglycans are particularly susceptible to Golgi dysfunction (Figure 5A – proteins in green). Collagen types IV, V and VI, fibronectin (FBN), and nidogen 2 (NID2) also showed reduced abundance in the media from both KO lines compared to WT (Figure 4A). Similarly, the secretion of matrix regulating enzymes such as FAM20C Golgi Associated Secretory Pathway Kinase (FAM20C) and PLOD2, as well as signalling molecules like TGFb, Growth Differentiation Factor 6 (GDF6) and LTBP2, was significantly downregulated in both KOs. Surprisingly, the protein showing the greatest increase in secretion in both KO lines was the intracellular protein Rho GDP Dissociation Inhibitor Beta (ARHGDIB), indicating aberrant translocation to the extracellular space.

As well as these similarities, we also observed many golgin-specific changes in this data set (Figure 5A - proteins circled in purple differentially impacted by each KO, Supplemental Figure 4C). For instance, thrombospondin secretion is more greatly affected in GMAP210 KO cultures, whilst Golgin-160 KO cells fail to efficiently secrete numerous additional ECM proteins and regulators like laminins and TIMP metalloproteases. Several components of the complement system also showed reduced secretion in the Golgin-160 KO cell lines implying this pathway may be impaired following Golgin-160 loss (Figure 5Aii and Supplemental Figure 4C). Overall, most proteins impacted by Golgin-160 loss showed reduced rather than increased abundance in culture media. In GMAP210 KO cells on the other hand, there were a significant number of proteins showing increased secretion, although these tended not to be ECM components. Of note, several lysosomal enzymes were released from GMAP210 KO cells suggesting either poor sorting at the Golgi or increased lysosome secretion (Figure 5Aii and Supplemental Figure 4C). In conclusion, although the secretion of a large subset of ECM proteins is broadly susceptible to loss of golgin function, there are also cargoes that show specificity with respect to their reliance on particular golgins.

### Golgin KOs induce changes in gene expression for some secreted proteins

Changes in secreted protein abundance in cell culture medium could arise due to altered gene expression of the cargos or altered trafficking to the cell surface. To test the former, we performed RT-PCR to measure the expression of cargo proteins identified in the secretome analysis. The proteins showing the greatest negative and positive fold change in abundance in the media of the KO cells were biglycan and ARHGDIB respectively. Consistent with this, we found that the expression of mRNA encoding each protein mirrored these changes, with biglycan expression reduced and ARHGDIB expression increased in KO clones (Figure 5B,C). To determine whether gene expression correlated with secretion more widely, three more targets were tested by RT-PCR: THBS1, which is only affected in GMAP210 KO cells; PLOD1, which is only impacted by Golgin-160 loss; and FAM20C, which shows reduced secretion in both KO lines. The expression of FAM20C was reduced in all KO clones, in line with secretion (Figure 5D). However, PLOD1 expression was not significantly altered in any of the KO lines (Figure 5E) and THBS1 expression was not consistently impacted (Figure 5F). Gene expression levels therefore did not fully predict secretion patterns for all targets, suggesting golgin loss can lead to changes to ECM secretion in different ways.

### Biglycan trafficking is unaffected by golgin loss

We next wanted to determine the direct impact of our golgin mutations on cargo transport and modification. For this we focused on biglycan as the protein showing the greatest fold change in secretion in all lines. Stable cell lines were generated expressing a construct encoding biglycan tagged with both mScarlet and a streptavidin binding protein (SBP) to allow live imaging of exogenous biglycan transport under the control of the RUSH trafficking system (Boncompain et al., 2021). In this system, tagged biglycan is expressed alongside a streptavidin-tagged ER hook, which binds the SBP tag on the biglycan and holds it in the ER. The addition of biotin prior to imaging is then used to outcompete this interaction and release biglycan for anterograde trafficking. In these experiments, the ER hook (not fluorescently labelled) is transiently expressed on a bi-cistronic vector that also encodes BFP-tagged Mannosidase-II (MannII-BFP) to provide a Golgi marker.

In the absence of ER hook, stably expressed BGN-SBP-mSc was localised to both the ER and Golgi in all cell lines (Supplemental Figure 5). After transient transfection with the ER hook, successful anchoring of the tagged biglycan was confirmed by localisation of fluorescence to the ER at steady state (Figure 6Ai, Bi, Ci T 00:00). Upon the addition of biotin, biglycan began to accumulate in cytoplasmic punctae, presumed to be ER exit sites, before moving towards the Golgi region in punctate carriers (Supplemental Movies 4-6, Figure 6Aii, Bii, Cii). Arrival at the Golgi region occurred approximately 14 minutes after biotin addition in all cases (Figure 6D). Here, the biglycan signal initially accumulated adjacent to the ManII-BFP labelled structures, but then showed increasing colocalization with ManII-BFP as it progressed through the stack (Supplemental Movies 4-6, Figure 6Ai, Bi, Ci). Finally, approximately 20 minutes after arrival at the Golgi, the emergence of post-Golgi carriers carrying biglycan to the cell periphery was readily observed (Figure 6Aiii, Biii, Ciii, E). Overall, transport kinetics were similar in all lines tested (Figure 6D,E), suggesting biglycan secretion is unimpeded by loss of GMAP210 or Golgin-160, at least in an over-expression bulk transport system. The changes in Golgi organisation observed in KO lines do not therefore significantly impact general cargo transport.

**Figure 6.**
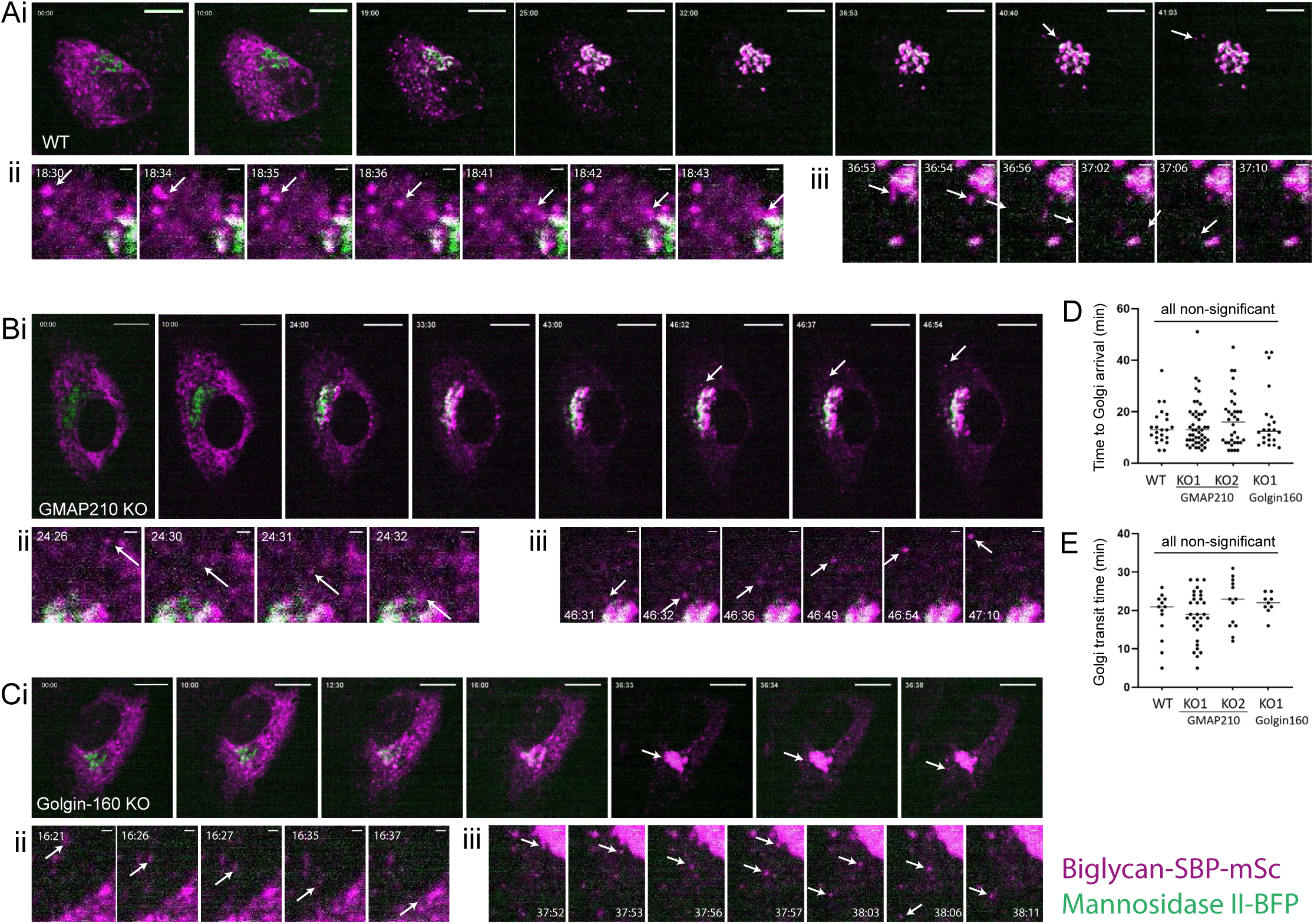
Biglycan can traffic efficiently in golgin KO cells. **A-D.** Biglycan RUSH assay in WT (A), GMAP210 KO (B) and Golgin-160 KO (C) cells stably expressing BGN-SBP-mSc (magenta) and transiently transfected with an ER hook (not visible) and Mannosidase II-BFP (green) prior to experiment. **A-C.** Single plane confocal images taken from time-lapse movies of biglycan transport after release from the ER by biotin addition at T 00:00. Time post biotin addition is indicated in top left corner as mm:ss. **Ai, Bi, Ci.** White arrows highlight the emergence of post-Golgi carriers. Scale bar 10 µm. **Aii, Bii, Ciii.** Crop showing incidence of ER-Golgi transport of biglycan as highlighted by arrow. Scale bar 1 µm. **Aiii, Biii, Ciii.** Crop showing incidence of post-Golgi transport of biglycan with carrier highlighted by arrow. Scale bar is 1 µm. **D.** Quantification of ER-Golgi transport, measured as time between biotin addition and the appearance of biglycan signal adjacent to Mannosidase II-BFP label. **E.** Quantification of Golgi transit time, measured as the time between biglycan enrichment adjacent to mannosidase II-BFP signal and the emergence of a visible post-Golgi carrier. **D-E.** All quantification performed on live movies as represented in A-C. Individual data points represent individual cells imaged across six independent experiments, bars show mean and standard deviation. **D.** Data were subjected to a Shapiro Wilk test for normality (failed) and then a nested one way ANOVA with Kruskall-Wallis test. **E.** Data were subjected to a Shapiro Wilk test for normality (passed) and then a nested one way ANOVA with Dunnet’s test for multiple comparisons to generate p values.

### Post-translational modification of biglycan is impaired in golgin KO cell lines

The inter-connectivity of Golgi cisternae and their spatial organisation is important to support and optimise the function of Golgi resident enzymes that post-translationally modify cargoes (Zhang and Wang, 2016). ECM proteins are often highly modified and these modifications are important to their assembly (Adams, 2023). We therefore sought to determine whether the disruption to Golgi organisation in our mutants impacts ECM protein modification, which may in turn contribute to assembly defects. Proteoglycans are heavily modified by the addition of large, sulphated GAG chains, making them highly susceptible to Golgi dysfunction. We therefore continued to focus on biglycan as a model cargo.

To look for changes in modification, cells stably expressing BGN-SBP-mSc were grown in serum free media overnight before collecting the media and lysate fractions of cultures for western blotting. Immunoblotting for both biglycan and the mScarlet tag showed that exogenous biglycan is secreted by all mutant lines (Figure 7A), consistent with the RUSH assays. Furthermore, the ratio of intracellular to extracellular protein was maintained, suggesting secretion rates are similar (Figure 7B). We did, however notice a shift in the molecular weight distribution of secreted biglycan in mutant cultures. In WT conditioned media, secreted BGN-SBP-mSc was identifiable as a broad band at around 250 kDa, with a small amount of protein running at 150kDa. In all KO lines however, a higher proportion of the protein ran at the lower molecular weight (Figure 7A). Calculation of the normalised ratio of 250 kDa vs 150 kDa protein confirmed this observation (Figure 7C). Blotting with the higher affinity RFP antibody also showed the presence of fully unmodified BGN-SBP-mSc at 75kDa in the media fraction, which was again secreted to a greater extent by KO cells (Figure 7A). These data suggest biglycan post-translational modification is perturbed in a similar way in the absence of either golgin.

**Figure 7.**
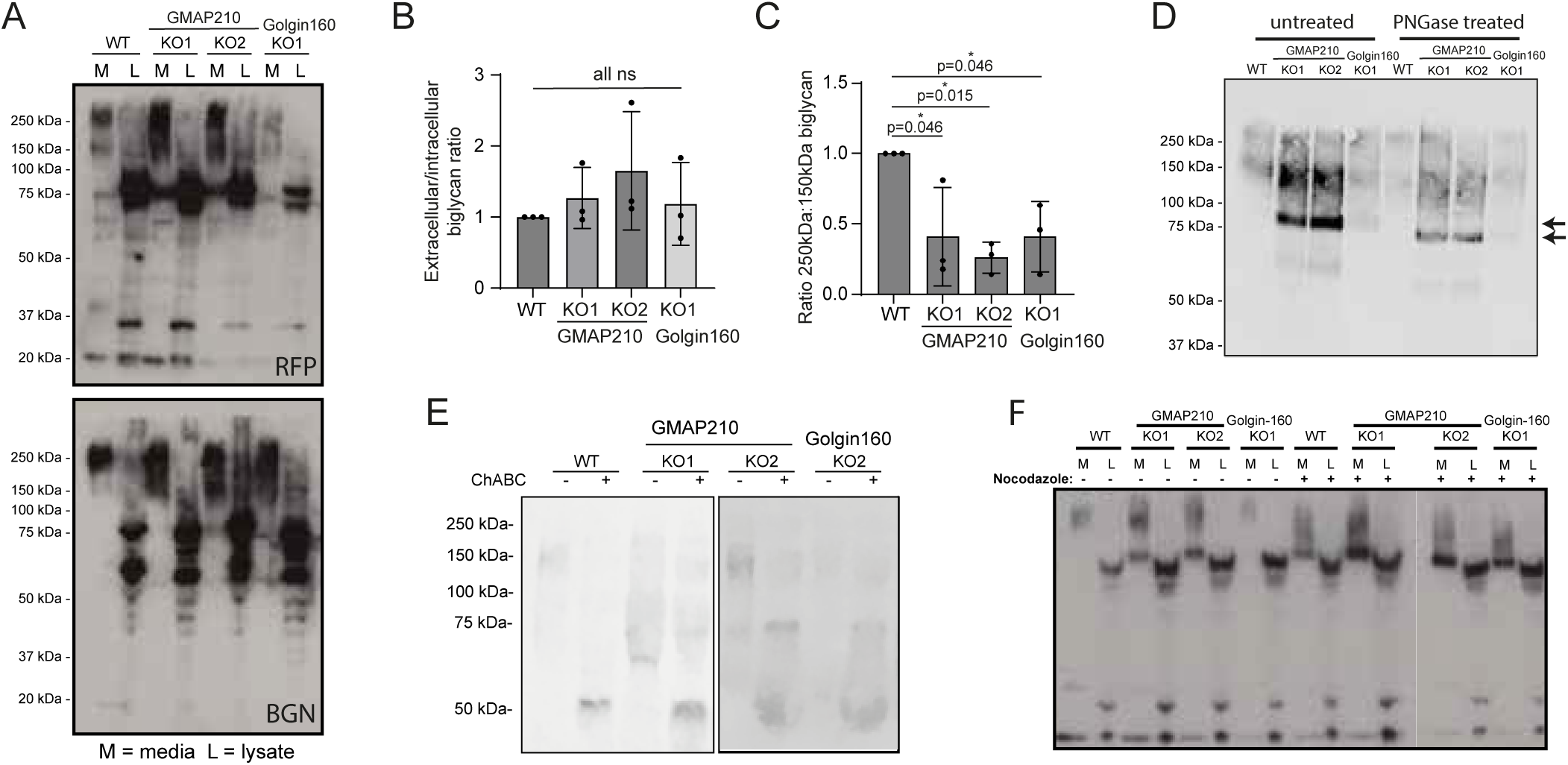
Biglycan is under-modified in golgin KO cells. **A.** Western blots of media (M) and lysate (L) samples taken from WT, GMAP210 KO and Golgin-160 KO cells stably expressing BGN-SBP-mSc, probed with anti-RFP and anti-biglycan antibodies. **B.** Quantification of the ratio between extracellular (M) and intracellular (L) biglycan in WT and KO cultures, as determined by densitometry of the blots shown in (A). **C.** Quantification of the ratio between 250kDa and 150kDa BGN bands present in the blots represented in (A). **B-C.** Dots represent measurements from independent experiments, bars show mean and SD. Data were subjected to a Shapiro Wilk test for normality (passed) and then a nested one way ANOVA with Dunnet’s test for multiple comparisons to generate p values. **D-E.** Western blots of medium and lysate samples from WT and KO lines stably expressing BGN-SBP-mSc. Collected samples were treated with PNGase F (D) or chondriotinase ABC (E) prior to SDS-PAGE. Blots probed with antibodies targeting the mSc tag. **D**. Arrows depict undigested (top) and PNGaseF digested (bottom) core protein. **F.** Western blots of medium and lysate samples from WT and KO lines stably expressing BGN-SBP-mSc. Media were collected 5 hours after the addition of DMSO (control) or nocodazole to cultures. Immunoblot for mSc tag (RFP).

**Table 1.**
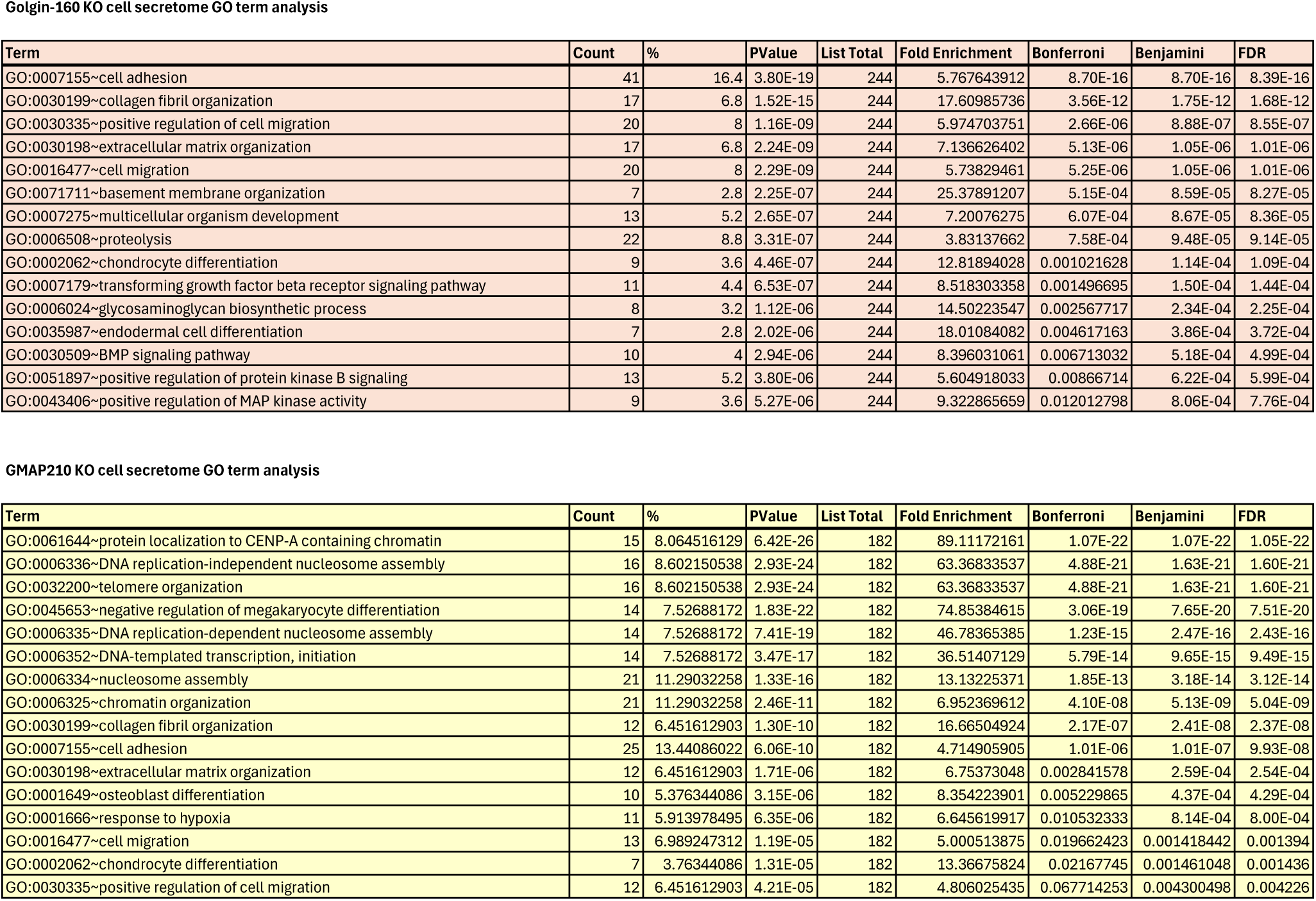
GO term analysis of golgin secretome data.

**Golgin-160 KO cell secretome GO term analysis**
| Term | Count | % | PValue | List Total | Fold Enrichment | Bonferroni | Benjamini | FDR |
| --- | --- | --- | --- | --- | --- | --- | --- | --- |
| GO:0007155~cell adhesion | 41 | 16.4 | 3.80E-19 | 244 | 5.767643912 | 8.70E-16 | 8.70E-16 | 8.39E-16 |
| GO:0030199~collagen fibril organization | 17 | 6.8 | 1.52E-15 | 244 | 17.60985736 | 3.56E-12 | 1.75E-12 | 1.68E-12 |
| GO:0030335~positive regulation of cell migration | 20 | 8 | 1.16E-09 | 244 | 5.974703751 | 2.66E-06 | 8.88E-07 | 8.55E-07 |
| GO:0030198~extracellular matrix organization | 17 | 6.8 | 2.24E-09 | 244 | 7.136626402 | 5.13E-06 | 1.05E-06 | 1.01E-06 |
| GO:0016477~cell migration | 20 | 8 | 2.29E-09 | 244 | 5.73829461 | 5.25E-06 | 1.05E-06 | 1.01E-06 |
| GO:0071711~basement membrane organization | 7 | 2.8 | 2.25E-07 | 244 | 25.37891207 | 5.15E-04 | 8.59E-05 | 8.27E-05 |
| GO:0007275~multicellular organism development | 13 | 5.2 | 2.65E-07 | 244 | 7.20076275 | 6.07E-04 | 8.67E-05 | 8.36E-05 |
| GO:0006508~proteolysis | 22 | 8.8 | 3.31E-07 | 244 | 3.83137662 | 7.58E-04 | 9.48E-05 | 9.14E-05 |
| GO:0002062~chondrocyte differentiation | 9 | 3.6 | 4.46E-07 | 244 | 12.81894028 | 0.001021628 | 1.14E-04 | 1.09E-04 |
| GO:0007179~transforming growth factor beta receptor signaling pathway | 11 | 4.4 | 6.53E-07 | 244 | 8.518303358 | 0.001496695 | 1.50E-04 | 1.44E-04 |
| GO:0006024~glycosaminoglycan biosynthetic process | 8 | 3.2 | 1.12E-06 | 244 | 14.50223547 | 0.002567717 | 2.34E-04 | 2.25E-04 |
| GO:0035987~endodermal cell differentiation | 7 | 2.8 | 2.02E-06 | 244 | 18.01084082 | 0.004617163 | 3.86E-04 | 3.72E-04 |
| GO:0030509~BMP signaling pathway | 10 | 4 | 2.94E-06 | 244 | 8.396031061 | 0.006713032 | 5.18E-04 | 4.99E-04 |
| GO:0051897~positive regulation of protein kinase B signaling | 13 | 5.2 | 3.80E-06 | 244 | 5.604918033 | 0.00866714 | 6.22E-04 | 5.99E-04 |
| GO:0043406~positive regulation of MAP kinase activity | 9 | 3.6 | 5.27E-06 | 244 | 9.322865659 | 0.012012798 | 8.06E-04 | 7.76E-04 |

**GMAP210 KO cell secretome GO term analysis**
| Term | Count | % | PValue | List Total | Fold Enrichment | Bonferroni | Benjamini | FDR |
| --- | --- | --- | --- | --- | --- | --- | --- | --- |
| GO:0061644~protein localization to CENP-A containing chromatin | 15 | 8.064516129 | 6.42E-26 | 182 | 89.11172161 | 1.07E-22 | 1.07E-22 | 1.05E-22 |
| GO:0006336~DNA replication-independent nucleosome assembly | 16 | 8.602150538 | 2.93E-24 | 182 | 63.36833537 | 4.88E-21 | 1.63E-21 | 1.60E-21 |
| GO:0032200~telomere organization | 16 | 8.602150538 | 2.93E-24 | 182 | 63.36833537 | 4.88E-21 | 1.63E-21 | 1.60E-21 |
| GO:0045653~negative regulation of megakaryocyte differentiation | 14 | 7.52688172 | 1.83E-22 | 182 | 74.85384615 | 3.06E-19 | 7.65E-20 | 7.51E-20 |
| GO:0006335~DNA replication-dependent nucleosome assembly | 14 | 7.52688172 | 7.41E-19 | 182 | 46.78365385 | 1.23E-15 | 2.47E-16 | 2.43E-16 |
| GO:0006352~DNA-templated transcription, initiation | 14 | 7.52688172 | 3.47E-17 | 182 | 36.51407129 | 5.79E-14 | 9.65E-15 | 9.49E-15 |
| GO:0006334~nucleosome assembly | 21 | 11.29032258 | 1.33E-16 | 182 | 13.13225371 | 1.85E-13 | 3.18E-14 | 3.12E-14 |
| GO:0006325~chromatin organization | 21 | 11.29032258 | 2.46E-11 | 182 | 6.952369612 | 4.10E-08 | 5.13E-09 | 5.04E-09 |
| GO:0030199~collagen fibril organization | 12 | 6.451612903 | 1.30E-10 | 182 | 16.66504924 | 2.17E-07 | 2.41E-08 | 2.37E-08 |
| GO:0007155~cell adhesion | 25 | 13.44086022 | 6.06E-10 | 182 | 4.714905905 | 1.01E-06 | 1.01E-07 | 9.93E-08 |
| GO:0030198~extracellular matrix organization | 12 | 6.451612903 | 1.71E-06 | 182 | 6.75373048 | 0.002841578 | 2.59E-04 | 2.54E-04 |
| GO:0001649~osteoblast differentiation | 10 | 5.376344086 | 3.15E-06 | 182 | 8.354223901 | 0.005229865 | 4.37E-04 | 4.29E-04 |
| GO:0001666~response to hypoxia | 11 | 5.913978495 | 6.35E-06 | 182 | 6.645619917 | 0.010532333 | 8.14E-04 | 8.00E-04 |
| GO:0016477~cell migration | 13 | 6.989247312 | 1.19E-05 | 182 | 5.000513875 | 0.019662423 | 0.001418442 | 0.001394 |
| GO:0002062~chondrocyte differentiation | 7 | 3.76344086 | 1.31E-05 | 182 | 13.36675824 | 0.02167745 | 0.001461048 | 0.001436 |
| GO:0030335~positive regulation of cell migration | 12 | 6.451612903 | 4.21E-05 | 182 | 4.806025435 | 0.067714253 | 0.004300498 | 0.004226 |

To better understand the nature of the biglycan modification defect, we first assessed N-glycosylation by subjecting secreted BGN-SBP-mSc to digestion by PNGase F, which cleaves N-glycan chains. After treatment, an identical small shift in biglycan molecular weight was observed in all samples indicating that this modification is intact in KO cultures (Figure 7D). Note, digestion had little impact on the higher molecular weight bands suggesting N-glycosylation does not contribute a significant amount to the mass of these biglycan forms. On the other hand, treatment of immunoprecipitated biglycan with chondroitinase ABC which digests GAG chains, did result in the loss of all higher and mid-molecular weight forms (Figure 7E). Interestingly, there was some resistance to digest in the GMAP210 KO clones that we attribute to altered modifications affecting enzyme-substrate affinity. Thus, we conclude that glycanation of biglycan is defective in the golgin KO cells.

One Golgi organisational feature common to both GMAP210 and Golgin-160 KO lines is fragmentation of the Golgi ribbon. To test if this would be sufficient to impair GAG chain synthesis, we fragmented the Golgi in our cell lines using the microtubule depolymerising agent nocodazole and repeated our secretion assays. Consistent with other studies (Harada et al., 2024), drug treatment impaired BGN modification in WT cells (Figure 7F). Surprisingly however, we also found that nocodazole treatment exacerbated the GAG modification defects in mutant cells, suggesting the observed fragmentation in KO lines was not fully pervasive. Overall, our studies show that GAG chain synthesis is impaired in the absence of both golgins tested.

## Discussion

In this study we show that loss of two *cis-*Golgi localised golgins causes distinct changes to Golgi morphology and function that ultimately lead to similar defects in ECM secretion, assembly and organisation. This is the first time that loss of Golgin-160 has been shown to impact ECM deposition and highlights the broad susceptibility of ECM cargoes to golgin loss. Altogether our data suggest that the impact of golgin depletion on ECM assembly is manifold, with gene expression, protein secretion, protein modification and ECM turnover all affected.

This study provides the most in-depth characterisation of Golgi structure in the absence of GMAP210 to date. In agreement with previous reports, we observed that loss of GMAP210 results in compaction of Golgi structures in the peri-nuclear region, fragmentation of the Golgi ribbon and cisternal dilation (Bird et al., 2018; Sato et al., 2015; Smits et al., 2010; Wehrle et al., 2019; Yamaguchi et al., 2021). We also note the concurrent unstacking of cisternae and build-up of tubulovesicular membranes in the peri-Golgi region. Altogether, this is indicative of a severe, general disruption to Golgi membrane dynamics. In 2014, Wong *et al* demonstrated that GMAP210 tethers GalNAcT2 containing vesicles and supports the correct localisation of the golgins giantin and GCC88 (Wong and Munro, 2014). Its loss may therefore impact enzyme sorting and transport directly, whilst indirectly triggering a cascade of problems in membrane trafficking by causing the mis-localisation of other Golgi organising proteins.

Loss of Golgin-160 did not seem to affect Golgi stacking or cisternal flattening, however there was a clear build-up of transport vesicles. Unfortunately, we were unable to confirm whether these were COPI vesicles at the light level as individual vesicles could not be distinguished in the peri-Golgi region. Regardless, the large number of vesicles in the peri-Golgi region in KO cells is consistent with a loss of vesicle tethering function, which may explain the large number of cargoes negatively impacted by Golgin-160 loss. A second feature of Golgi organisation in Golgin-160 KO cells was fragmentation of the Golgi ribbon. It has been reported that Golgin-160 recruits dynein to Golgi membranes (Yadav et al., 2012). Microtubules are required to support lateral tethering of cisternae, and thus reduced motor and microtubule recruitment in the absence of Golgin-160 may underly this phenotype. Note, we did not observe the Golgi positioning or anterograde transport defects reported previously in another system (Yadav et al., 2012).

In humans, severe and hypomorphic mutations in the Trip11 gene encoding GMAP210 cause achondrogenesis type 1A (ACG1A) and odontochondrodysplasia (OCDC) respectively, (Costantini et al., 2021; Del Pino et al., 2021; Medina et al., 2020; Qian et al., 2021; Upadhyai et al., 2021; Wehrle et al., 2019). These diseases are characterised by skeletal abnormalities such as shortened ribs and cranial-facial malformations. Study of *Trip11* KO mouse models has ascertained that a primary driver of pathology is defective ECM secretion during skeletal development (Bird et al., 2018; Follit et al., 2008; Smits et al., 2010; Yamaguchi et al., 2021). Here we report a comparable attenuation of ECM secretion in our GMAP210 KO epithelial lines, with many of our phenotypes recapitulating those of *in vivo* and patient studies, including impaired collagen fibrillogenesis (Yamaguchi et al., 2021) and diminished production of highly glycanated proteoglycans (Wehrle et al., 2019). Interesting, skeletal dysplasia has not been reported in existing studies of *Golga3/Golgin-160* mutant mice (Banu et al., 2002; Bentson et al., 2013; Matsukuma et al., 1999), which seems incongruous with the similarities seen here between GMAP210 and Golgin-160 KO ECM defects. Compensatory mechanisms often mask severe phenotypes in golgin mutant animal models, which may explain this discrepancy (Bergen et al., 2017). Whether more subtle *in vivo* phenotypes, such as altered bone mass density will be found with more extensive investigation remains to be determined.

Golgin-160 negatively impacted the secretion of a greater number of ECM components than GMAP210, including several laminins and metalloproteases. This implies it has a greater impact on transport. Consistent with this, Golgin-160 has been shown to bind to the Golgi-associated PDZ domain containing protein PIST/GOPC (Hicks and Machamer, 2005), which supports transport of cargoes to the plasma membrane. More specifically, the anterograde trafficking of the membrane-bound GLUT4 transporter (Williams et al., 2006) and β4 adrenergic receptor (Hicks et al., 2006) has been shown to require Golgin-160. Our results expand the list of golgin-160 dependent cargoes in the context of soluble protein secretion and show that extracellular matrix components are particularly affected.

Overall, whilst Golgin-160 KO had a primarily negative impact on general secretion, more than half of the cargoes impacted by GMAP210 loss showed increased secretion in KO cells compared to WT. Interestingly this was not the case for ECM proteins, which were almost always negatively impacted. One family of proteins specifically impacted by GMAP210 KO were the lysosomal enzymes cathepsins, four of which were aberrantly secreted by KO cells. This raises the possibility that the increased abundance of certain proteins in GMAP210 KO culture medium may stem from sorting defects. Interestingly, secreted cathepsins have been implicated in ECM degradation (Vidak et al., 2019; Vizovisek et al., 2019). Of note, two of those identified here (cathepsins B + K) have been shown to cleave targets such as SPARC (Podgorski et al., 2009), collagen type IV (Guinec et al., 1993), and fibronectin (Guinec et al., 1993), which were all reduced in abundance in the same KO media. Whether secreted cathepsins are contributing to the ECM phenotypes observed here therefore remains an intriguing open question, especially considering their potential as drug targets if found to contribute to disease pathology (Jangra et al., 2024).

Proteoglycans appear particularly susceptible to the loss of both GMAP210 and Golgin-160, with KO affecting both gene expression and glycanation of biglycan. The biosynthetic pathway for GAG chains serves as an excellent example of the importance of Golgi organisation (Hellicar et al., 2022; Ricard-Blum et al., 2024). Chains are built by enzymes organised sequentially across the Golgi stack and competition between enzymes determines which types of GAG chains are built. Consequently, chain chemistry is determined by relative enzyme abundance, enzyme distribution, and trafficking speeds, making it particularly sensitive to a variety of Golgi perturbations (Adusumalli et al., 2021; Ahat et al., 2022; Chang et al., 2013). In our case, a common structural feature in both GMAP210 and Golgin-160 KO lines is Golgi fragmentation. Previously, this has been shown to disrupt the balance of GAG chain species (Ahat et al., 2022; Harada et al., 2024) and reduce overall GAG sulphation (Ahat et al., 2022). We also find that nocodazole-induced Golgi fragmentation in RPE1 cells impairs biglycan GAG modification. Interestingly, nocodazole treatment of our mutant lines exacerbated glycan defects, suggesting that Golgi fragmentation is not fully pervasive. It is therefore worth considering the role of other golgin-specific changes to Golgi morphology.

The most prominent feature of Golgin-160 KO cells was the accumulation of transport vesicles. Loss of the COPI-binding COG complex has previously been shown to interfere with GAG chain length (Adusumalli et al., 2021), presumably due to impaired enzyme recycling. Thus, disruption to vesicular transport may contribute to glycan assembly defects in Golgin-160 KO cells. Meanwhile, a striking feature of GMAP210 KO cells was cisternal dilation, a feature also seen following the treatment of cells with actin destabilising drugs (Lazaro-Dieguez et al., 2006). One such drug, cytochalasin B, also inhibits proteoglycan synthesis without impacting secretion (Lohmander et al., 1979). Thus, cisternal dilation may contribute to glycan under-modification in the KO cells, perhaps by impacting the frequency with which membrane-bound enzymes interact with soluble luminal cargo and so reducing the efficiency of modification. Overall, it seems likely that the combined effect of different perturbations to Golgi homeostasis in each mutant cell line contribute to impaired glycanation.

In conclusion, we propose that golgins have distinct, non-redundant functions but work collectively to create the correct physical and biochemical space to support efficient ECM protein secretion and modification.

---

**Supplemental Table 1 GO term analysis of golgin Secretome data**

Proteins that showed significantly altered abundance in KO cell derived media compared to WT conditioned media (p value < 0.01) were entered into the DAVID knowledgebase (https://david.ncifcrf.gov/tools.jsp) and analysed against the Biological pathways GO-BP-DIRECT list for Gene Ontology term enrichment. The 15 GO terms showing the most significant enrichment, as determined by p-value, are presented here.

**Supplementary Figure 1.**
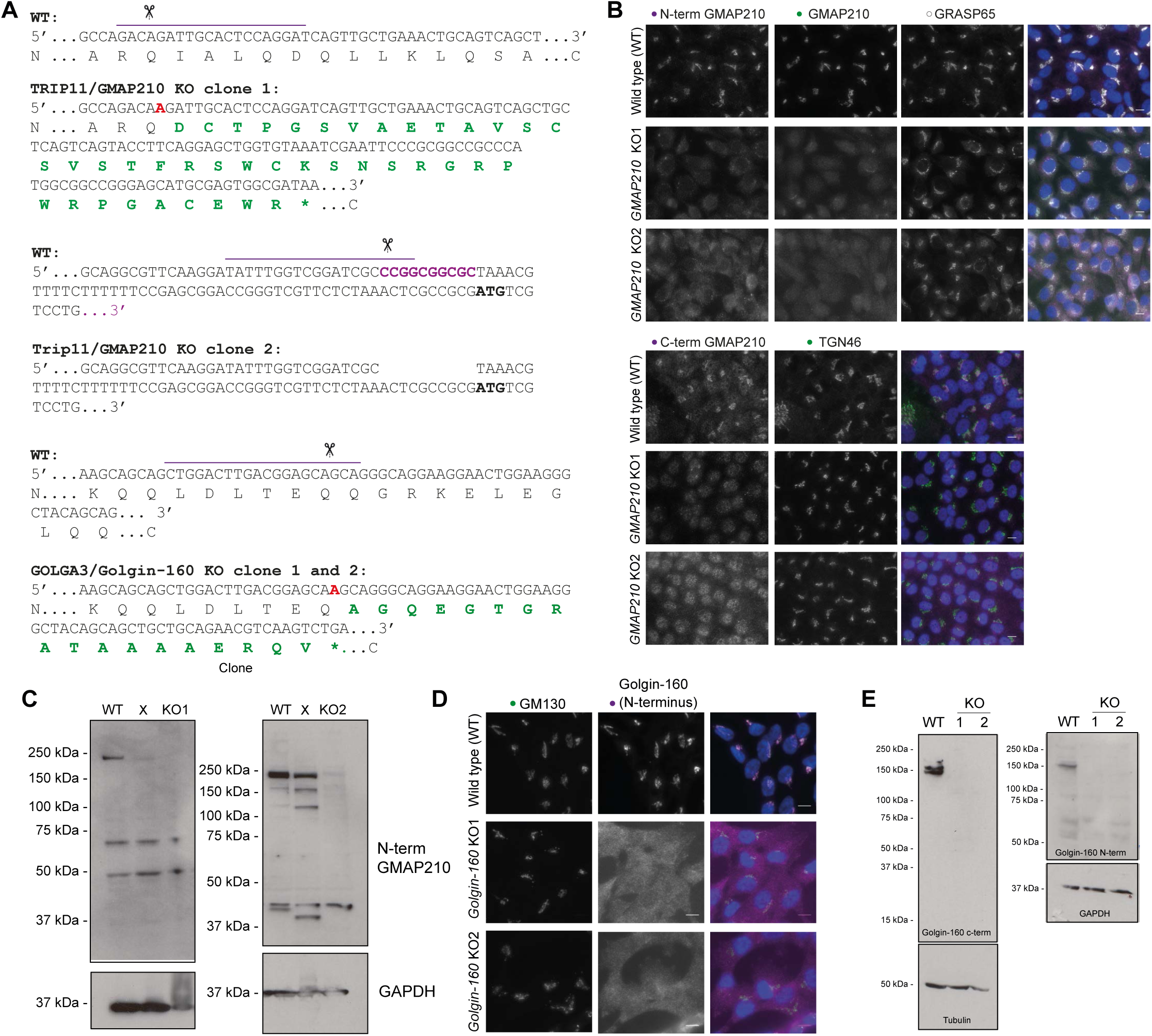
Generation of CRISPR-Cas9 mutant lines. **A.** CRISPR design and resulting mutations in chosen knockout clones. Top lines show gene sequence in wildtype (WT) and mutant (KO) clones with encoded amino acid sequence underneath. Purple lines indicate gRNA target sequence, scissors point to predicted Cas9 cut site, red letters in KO sequences indicate mutagenic base pair insertions, purple letters in WT sequence indicate base pairs deleted in the mutant lines. Green amino acids are mutagenic changes arising after frameshift in KO lines and * denotes a premature stop codon. **B.** Maximum projection widefield images of WT and GMAP210 KO lines immunolabelled as indicated. GMAP210 antibodies were raised against amino acids 14-148 (N-term GMAP210), 159-365 (GMAP210), and 1760-1855 (C-term GMAP210). **C.** Western blots of WT and GMAP210 KO cells probed with an antibody targeting amino acids 14-148 or GMAP210 (central well is an unsuccessful clone - X). **D.** Maximum projection widefield images of WT and Golgin-160 KO clones labelled with GM130 (green, *cis-*Golgi) and Golgin-160 (magenta). **D.** Western blot analysis of WT and Golgin-160 KO cell lysates probed with Golgin-160 antibodies and tubulin and GAPDH as loading controls. **D, E.** Golgin-160 N-terminal and C-terminal antibodies raised against amino acids 1-350 and 1436-1498 respectively.

**Supplemental Figure 2.**
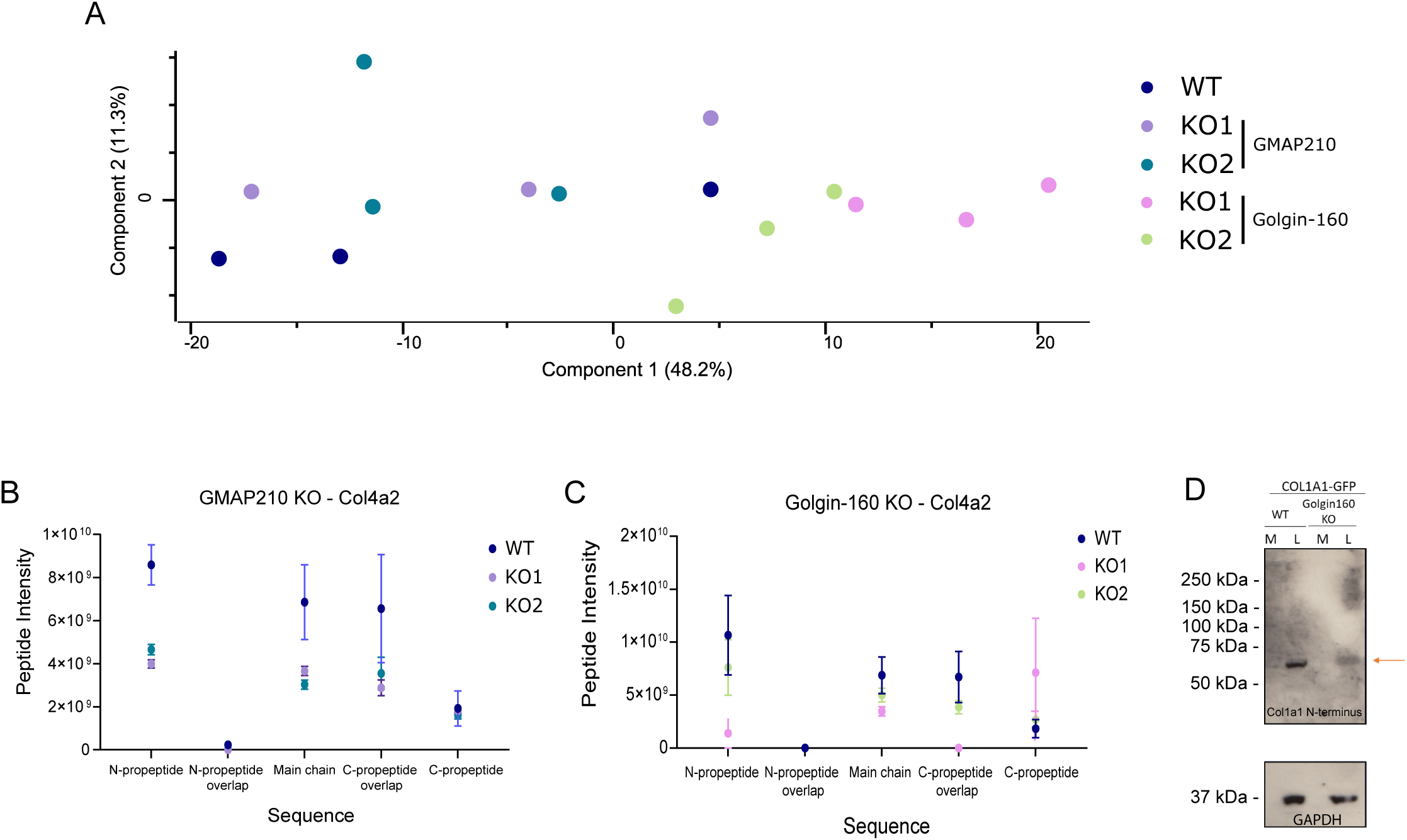
Matrisome analysis in mutant cells. **A**. Principal component analysis of the mass spectrometry experiment shown in Figure 3. **B, C.** The Col4a2 sequence was split into its structural features, abundances of relevant peptides were averaged within each feature and plotted for each mutant. Total Col4a2 abundance was normalised across conditions to investigate peptide level variation. N=3. **D.** Western blots of media (M) and lysate (L) fractions from WT and Golgin-160 KO cell cultures stably expressing pro-SBP-GFP-COL1A1. Blots probed with the LF39 antibody targeting the N-terminal propeptide domain of procollagen type I and GAPDH as a housekeeping protein.

**Supplementary Figure 3.**
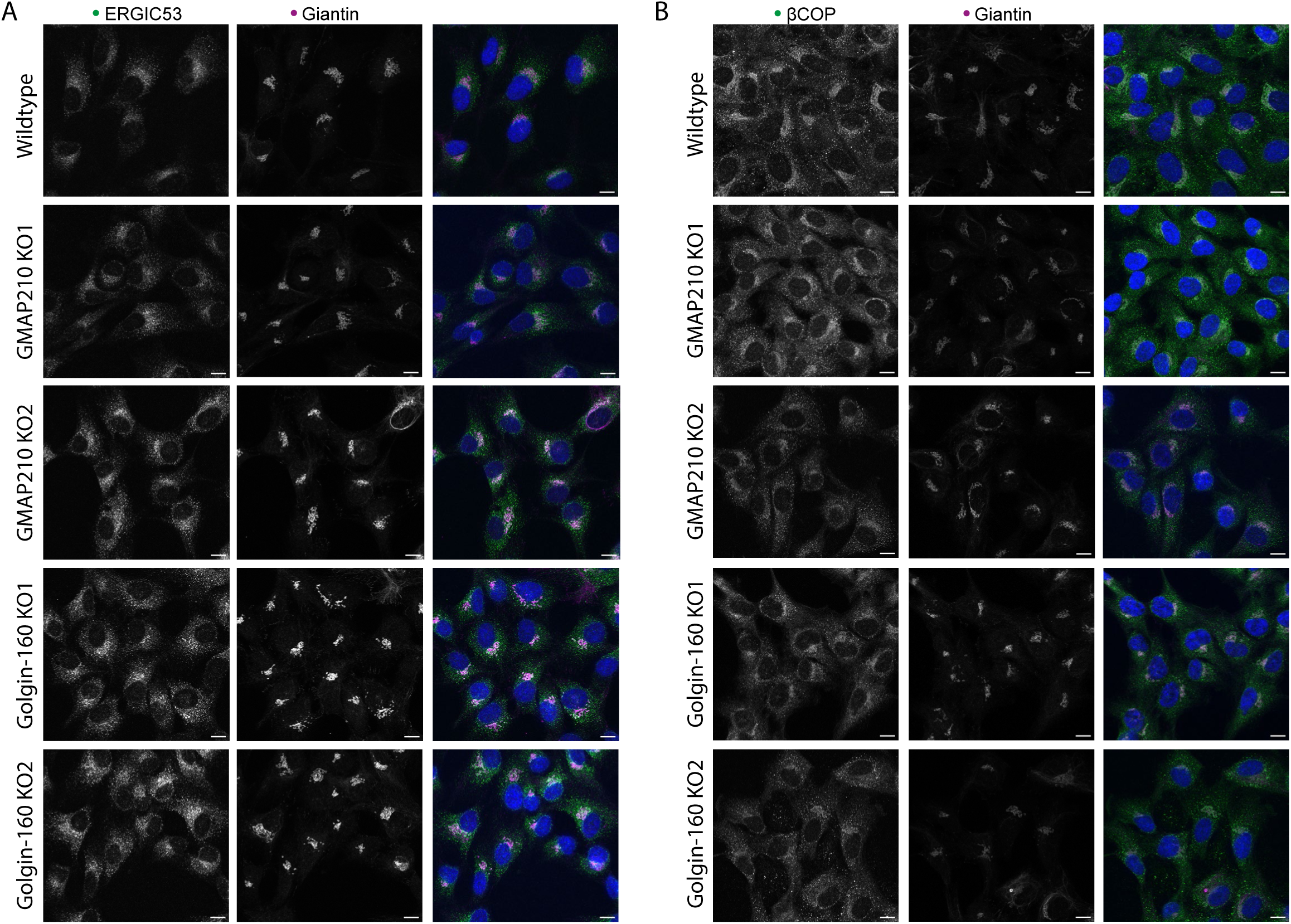
Early secretory pathway organisation in golgin mutants. **A-B.** Confocal maximum projection images of WT and golgin KO cell lines stained for cis/medial Golgi membranes (giantin, magenta) markers and either ERGIC (ERGIC53, green) (A) or COP1 (βCOP, green) structures (B). Scale bars 10 µM.

**Supplemental Figure 4.**
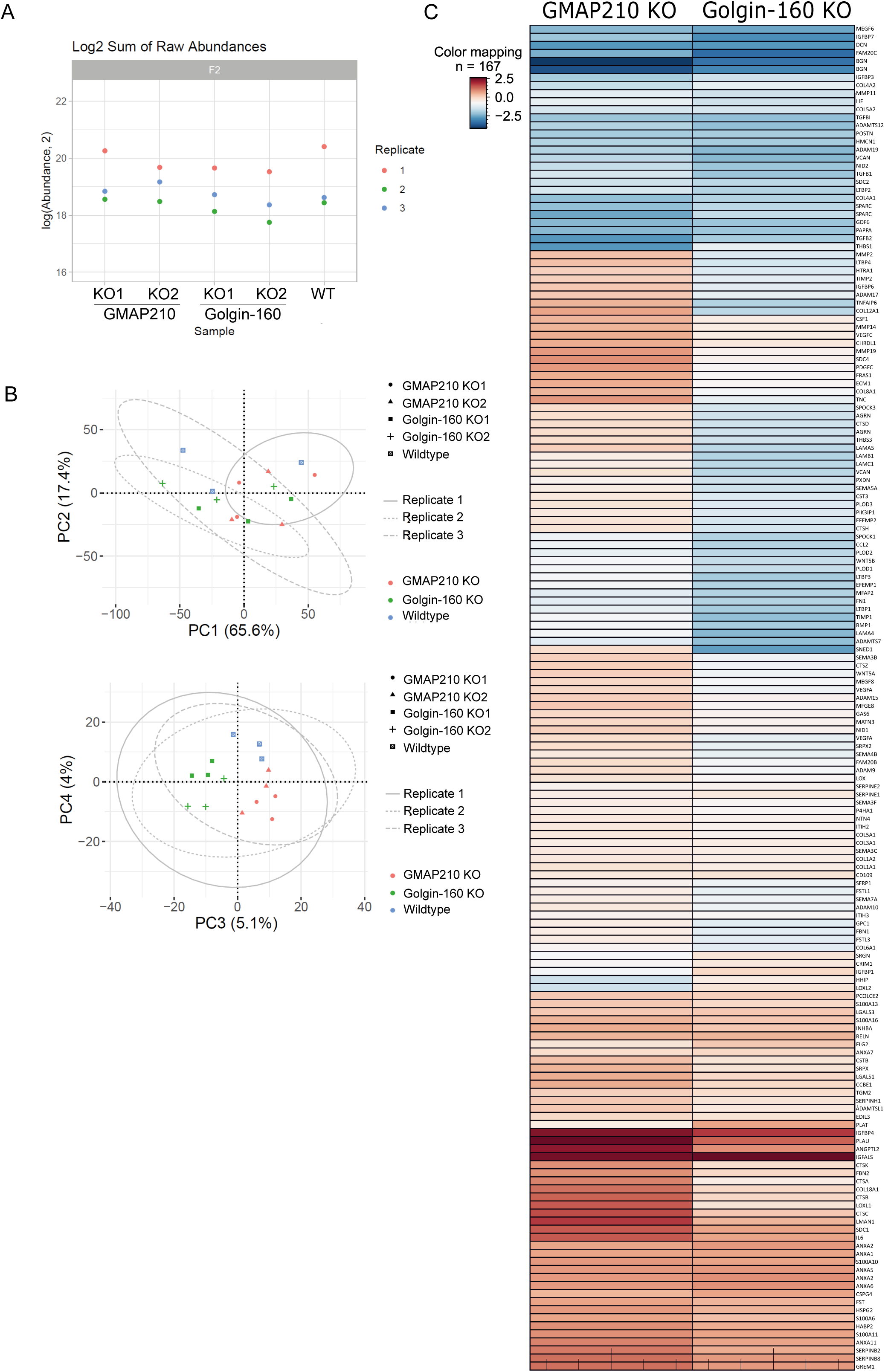
Secretome analysis in mutant cells. **A.** Log2 sum of raw peptide abundance in each proteomics replicate from the secretome analysis shown in Figure 5. **B.** Principal component analysis of the secretome data sets represented in Figure 5. **C.** Heat map comparing protein abundance changes between the two different golgin KOs in the secretome experiment. Red and blue indicate increased and decreased abundance respectively, intensity of colour is determined by average log-fold change across mutants compared to WT.

**Supplementary Figure 5.**
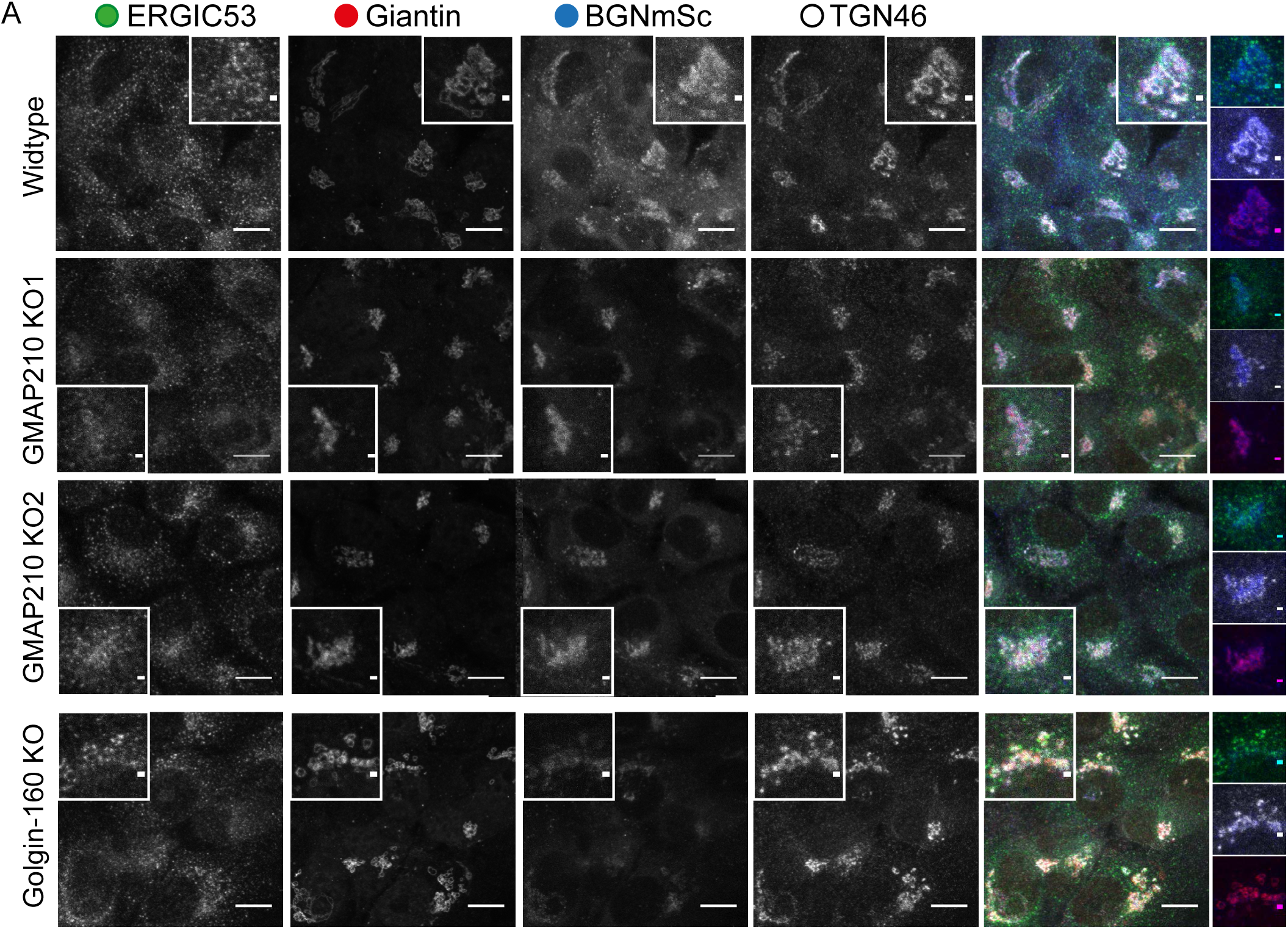
BGN-SBP-mSc localisation in mutant cells. **A.** Widefield maximum projections of cells stable expressing BGN-SBP-mSc (blue) and immunolabelled for the ERGIC (ERGIC53, green), cis/medial Golgi (giantin, magenta) and TGN (TGN46, greyscale). Scale bar in main image 10 µm and insert 1 µm.

**Supplemental Movie 1 – Golgi reconstruction and segmentation in WT cell**

Electron tomographic reconstruction of a 250 nm section through a WT cell showing an organised interconnected Golgi ribbon. Segmentation of a section of this ribbon shows cisternal membranes (labelled in blue/purple), dilated structures at the TGN (red), tubulovesicular structures (yellow), and vesicles (green). Movie shows slices running top-bottom-top through the section, then top to bottom adding the segmentation, followed by a zoom and rotational views of the 3D segmented structures. Movie runs at 40 frames per second.

**Supplemental Movie 2 – Golgi reconstruction and segmentation in GMAP210 KO2 cell**

Electron tomographic reconstruction of a 250 nm section through a GMAP210 KO2 cell showing scattered and dilated Golgi structures. Segmentation of two closely apposed Golgi stacks shows cisternal membranes (labelled in blue/purple), dilated structures (red), tubulovesicular structures (yellow), and vesicles (green). Movie shows slices running top-bottom-top through the section, then top to bottom adding the segmentation, followed by a zoom and rotational views of the 3D segmented structures. Movie runs at 40 frames per second.

**Supplemental Movie 3 – Golgi reconstruction and segmentation in Golgin-160 KO2 cell**

Electron tomographic reconstruction of a 250 nm section through a Golgin-160 KO2 cell showing a fragmented Golgi, with stacks surrounded by vesicles. Segmentation of one stack shows cisternal membranes (labelled in blue/purple), dilated structures at the TGN (red), tubulovesicular structures (yellow), and vesicles (green). Movie shows slices running top-bottom-top through the section, then top to bottom adding the segmentation, then a zoom and rotational views of the 3D segmented structures. Movie runs at 40 frames per second.

**Supplemental Movie 4 – Biglycan RUSH in WT cell**

Spinning disc confocal timelapse movie of a biglycan RUSH assay in a WT cell. Cells are stably expressing BGN-SBP-mSc (magenta) and transiently expressing KDEL-streptavidin (unlabelled) and Mannosidase-II-BFP (green). Movie begins 15 minutes after biotin addition (time after biotin indicated in top left (mm:ss). One frame taken every 500 ms. Scale bar 10 µm.

**Supplemental Movie 5 – Biglycan RUSH in GMAP210 KO2 cell**

Spinning disc confocal timelapse movie of a biglycan RUSH assay in a GMAP210 KO2 cell. Cells are stably expressing BGN-SBP-mSc (magenta) and transiently expressing KDEL-streptavidin (unlabelled) and Mannosidase-II-BFP (green). Movie begins 15 minutes after biotin addition (time after biotin indicated in top left (mm:ss). One frame taken every 500 ms. Scale bar 10 µm.

**Supplemental Movie 6 – Biglycan RUSH in Golgin-160 KO2 cell**

Spinning disc confocal timelapse movie of a biglycan RUSH assay in a Golgin-160 KO2 cell. Cells are stably expressing BGN-SBP-mSc (magenta) and transiently expressing KDEL-streptavidin (unlabelled) and Mannosidase-II-BFP (green). Movie begins 10 minutes after biotin addition (time after biotin indicated in top left (mm:ss). One frame taken every 500 ms. Scale bar 10 µm.

## Methods

### Cell culture

Telomerase immortalised human retinal pigment epithelial cells (hTERT-RPE1) were purchased from the American Type Culture Collection (ATCC) and grown in DMEM-F12-HAM (Life technologies #11320-033) supplemented with 10% decomplemented fetal calf serum (FCS, Gibco A5256701), passaging 1:10 every 3-4 days. HEK293T cells used for making lentivirus were grown in DMEM (D5796) supplemented with 10% FCS. Transient transfections of RPE1 cells were performed using Lipofectamine 2000 (Invitrogen catalog #11668027) and 2 µg DNA per 35 mm dish according to the manufacturer’s instructions. Stable cell lines were generated using Lenti-X^TM^ Packaging Single Shots kit (Takara, #631275) according to kit instructions and as described in the next section.

### Generating knockout cell lines

#### Transfection with RNPs

To generate GMAP210 KO lines the IDT ALT-R™ CRISPR system was used according to the manufacturer’s protocols. Two crRNAs with sequences ATCCTGGAGTGCAATCTGTC (designed by IDT and targeting exon 4) and TATTTGGTCGGATCGCCCGG (designed by Benchling targeting exon 1) were ordered from IDT. To duplex the gRNAs, 1 µl of 1 nmol/µl Alt-R™ CRISPR-Cas9 crRNA and 1 µl of 1 nmol/µl Alt-R™ CRISPR-Cas9 tracrRNA (IDT #1072533) were mixed with 98 µl duplex buffer (IDT #1072570) and heated to 95°C for 5 minutes before cooling on the bench. Alt-R™ S.p. Cas9 Nuclease V3 (IDT #10010588 LOT 000088243) was diluted to 1 µM in Cas9 working buffer (20 nM HEPES, 150 nM KCl pH 7.5). To make the RNP, 1.5 µl duplexed gRNA + 1.5 µl 1 µM Cas9 + 0.6 µl Cas9 PLUS reagent (Invitrogen™ Lipofectamine™ CRISPRMAX™ #CMAX00001) + 21.4 µl optiMEM were mixed per transfection. RNP mixes were left for 5 minutes at room temperature then 25 µl RNP mix + 1.2 µl CRISPRMAX (Invitrogen™ Lipofectamine™ CRISPRMAX™ #CMAX00001) + 23.8 µl of optiMEM (Gibco #31985-070) were added to a well of a 96 well plate. This was left to incubate for 20 minutes at room temperature. Meanwhile RPE1 cells were trypsinised, counted, centrifuged at 1000 xg for 3 minutes and then resuspended at a concentration of 20000 cells per 100 µl. After the 20- minute transfection incubation, 100 µl cell suspension (20000 cells) was added to the wells and plates put at 37°C 5% CO2. Cells were then expanded, and successful gene knockout was determined by immunofluorescence staining of transfected cells.

#### Transfection with lenti-CRISPR

Golgin-160 KO lines were generated using lentiCRISPRv2 constructs. To make the construct, two single stranded oligos targeting exon 20 of the GOLGA3 gene (transcript GOLGA3-101 on ensemble) plus vector overhangs were ordered from IDT. Sequences: F1: 5’ CACCGCTGGACTTGACGGAGCAGCA 3’ R1: 5’ AAACTGCTGCTCCGTCAAGTCCAGC 3’ (red is gene targeting). Oligos were annealed in T4 ligation buffer (NEB #B0202A) with T4 PNK (NEB M0201S) for 30 minutes at 37°C followed by a 5-minute incubation at 95°C and cooling by ramping down to 25°C at 5°C/minute. Annealed oligos were ligated into the lentiCRISPRv2 vector (Addgene Plasmid #52961) overnight using T4 ligation buffer (NEB #B0202A) and T4 ligase (NEB #M0202S). Ligations were transformed into One Shot™ Stbl3™ Chemically Competent *E. coli* (Invitrogen™ #C737303) according to the manufacturer’s instructions and bacterial cultures grown on ampicillin supplemented agar plates (BD #214530). Successful ligation was determined by extracting constructs with a QIAprep spin mini-prep kit (QIAGEN #27106) from bacterial colonies, followed by sequencing (MWG Eurofin). The plasmid was then transfected into HEK293T cells using the Lenti-X^TM^ Packaging Single Shots kit (Takara Bio # 631278) according to kit instructions. After 48 hours, virus was harvested and 8 µg/ml polybrene (Santa Cruz Technology sc-134220) added to a 1 ml aliquot. Media was aspirated from a 6 cm dish of RPE1 cells and the virus aliquot added to cells for incubation for 1 hour at 37°C. DMEM-F12-HAM supplemented with 8 µg/ml polybrene was then added to cells and cultures were incubated for 48 hours at 37°C. Transfected cells were selected at the next passage by supplementing media with 10 µg/ml puromycin dihydrochloride (Santa-cruz technologies sc-1080701). Successful gene editing was determined by immunofluorescence staining of transfected cells.

#### Making clones

To identify gene edited cells, CRISPR/Cas9 transfected cell lines were single cell sorted into a 96 well dish. Clones were expanded and KO confirmed by immunofluorescence and western blot. DNA was extracted from confirmed KO clones using a PureLink genomic DNA mini kit (Invitrogen #K1820-02) and the CRISPR target region was amplified by PCR using Q5® Hot Start High-fidelity 2x master mix (NEB #M0493L) according to manufacturer’s instructions and touch down PCR programme: 1) 95°C for 3 minutes, 2) melt at 95°C for 25 seconds, 3) anneal at 72°C for 25 seconds, 4) extension at 72°C for 45 seconds, 5) repeat steps 2-4 reducing the annealing temperature by 0.5°C per cycle x 20 cycles, 6) melt at 95°C for 25 seconds, 7) anneal at 72°C for 25 seconds, 8) extension at 72°C for 45 seconds 9) repeat steps 6-8 x25 10) 72°C 5 minutes 11) 4°C hold.

Golgin-160/GOLGA3 genotyping primers: Fwd 5’TGGGAGGTGGATCAGAAAGA3’ Rev 5’ACCCTCCCATTTGCTGTAGT3’. GMAP210/TRIP11 genotyping primers for exon 4: Fwd 5’ GTTGATAGAGGACCATCATTGGA 3’ Rev 5’ TACACCAGCTCCTGAAGGTA 3’. GMAP210/TRIP11 primers for exon 1: Fwd 5’TTCGTGGGAATGAGCAGAAG3’ Rev 5’ CAAGAGGAGGTGTGTGAAGAAA3’.

PCR products were purified using Qiaquick PCR purification kit (QIAGEN #28104) and A-tailed by mixing 3 µl PCR product + 1 µl 10x Thermopol buffer (NEB M0267S) + 0.2 µl ATP + 1 µl Taq polymerase and incubating the reaction at 70°C for 30 minutes. This reaction was then ligated into pGEM t-easy vector with T4 ligase and T4 ligation buffer (Promega #A1360) so that individual alleles could be sequenced. Sequencing was performed by MWG Eurofins using their stock T7 forward primer.

### Cloning and constructs

To make BGN-SBP-mSc, two gene blocks were designed using Hifi NEBuilder to encode the required tagged biglycan protein plus accompanying flanking sequences for sequential assembly into the pLVXNeo vector (Takara Bio #632181) using the NEB HiFi assembly system. The tagged biglycan coding region consisted of the biglycan mRNA sequence (transcript ID: <u>ENST00000331595.9</u>, CCDS: <u>CCDS14721</u>) followed by linker GSAGSAAGSGEF (Waldo et al., 1999), the streptavidin binding protein sequence, linker `GGGGSGGGGS`, and the mScarlet-i coding sequence. pLVXNeo was digested with XhoI (NEB #R0146S) in NEB 3.1 10x buffer (NEB #B7203S) for 15 minutes at 37C and then BsmBI- V2 (NEB #R0739S) was added for a second digest at 55°C for 20 minutes. Reactions were heated to 80°C for 20 minutes to inactivate enzymes. Gene block 1 was assembled into the digested vector with the Hifi assembly mix (NEB #E2621), and then the plasmid was transformed into NEB 5-alpha cells (C2987H), both according to manufacturer’s instructions. Colonies were grown up in LB plus ampicillin and plasmids were extracted using a QIAprep spin mini-prep kit (QIAGEN #27106). Successful assembly of insert into vector was confirmed by test digest with BsrGI and BamHI and sequencing. This intermediate vector was then digested with XhoI and assembled with the second block and confirmed as before.

Design of StrKDEL-IRES-mannosidase II-mTagBFP2 is detailed in (McCaughey et al., 2021) and available on Addgene (Plasmid #165460). Design of the pro-col1a1-SBP-GFP plasmid is detailed in (McCaughey et al., 2019) and available on Addgene (Plasmid #110726).

### Preparation of the cell derived matrix

To collect the cell derived matrix, cells were grown to confluence (either on coverslips or plastic) and their media supplemented with 50 μg/ml L-ascorbic acid-2-phosphate (Sigma-Aldrich A8960) for a further 7 days. Cells were then washed in PBS and the cell layer removed by incubation with pre-warmed extraction buffer (20 mM NH_4_OH (Sigma-Aldricth #221228) and 0.5% Triton X-100) for three minutes at room temperature. The dish was then washed three times with dH_2_O and incubated with 10 μg/ml DNase I (Roche) diluted in reaction buffer (10 mM Tris-HCl, pH 7.6, 2.5 mM MgCl2, and 0.5 mM CaCl2) for 30 min at 37°C. After a further 3 washes in water, the cell derived matrix was either fixed for staining or AFM imaging, or collected in sample buffer for assay (see appropriate section).

### Primary Antibodies

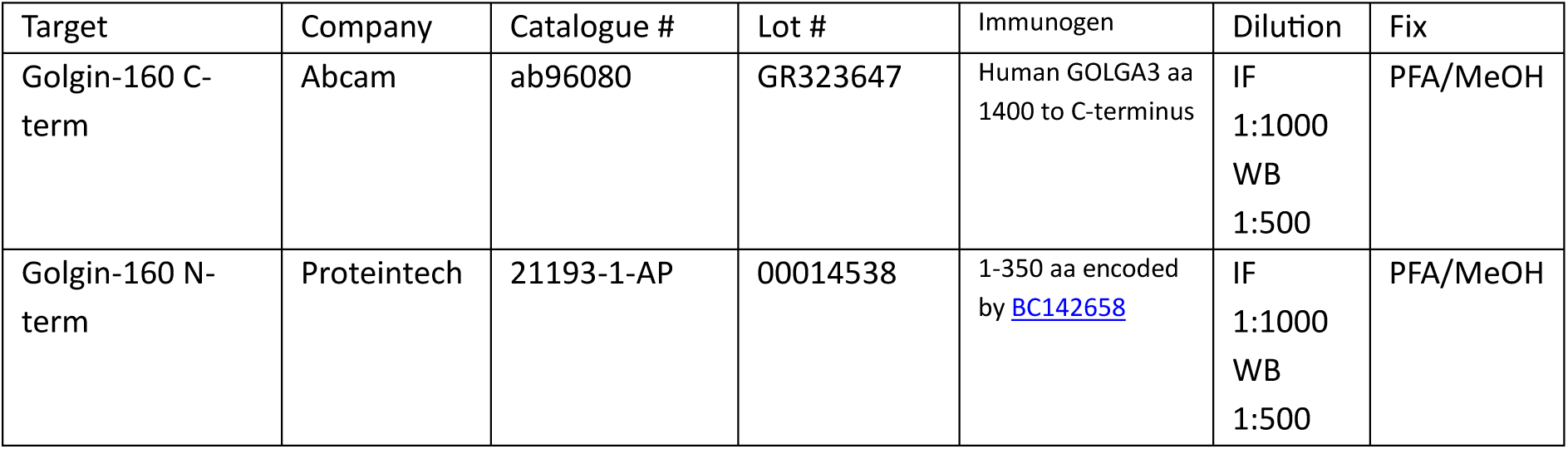
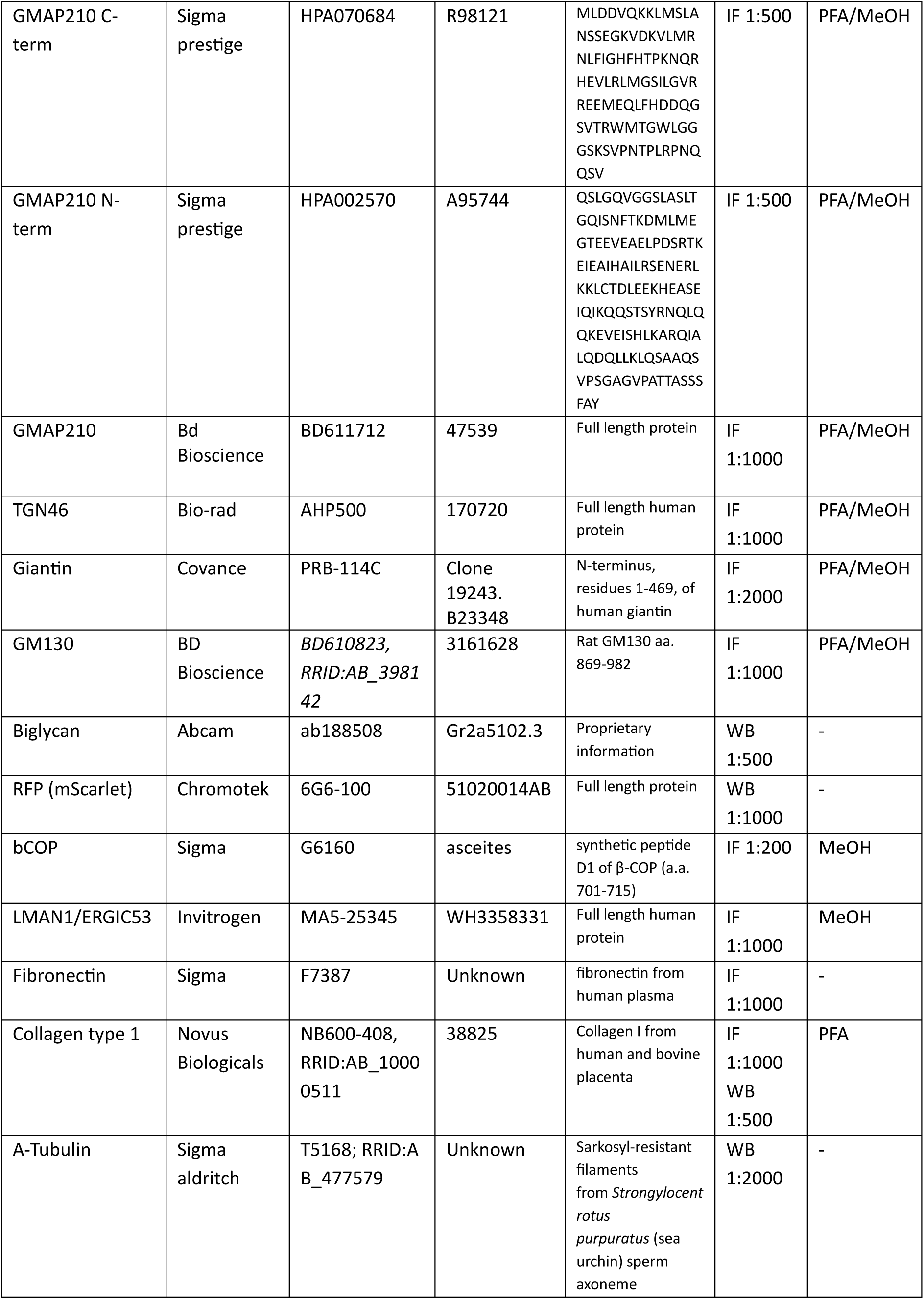

### Immunofluorescence and fixed cell microscopy

For antibody labelling, cells grown on autoclaved coverslips (Menzel #1.5, Fisher Scientific) were fixed with 4% PFA (Fisher-Scientific P/0840/53) for 10 minutes followed by permeabilisation for 10 minutes with 0.1% Tx100 (Sigma-Aldritch #X100) at room temperature. For ER antibodies, cells were fixed with MeOH (Sigma-Aldritch #32213) for 3 minutes at −20°C. Cells were then blocked with 3% BSA (Sigma-Aldritch #A9647) for 30 minutes before incubation with primary antibody for 30 minutes, three washes with PBS, and incubation with Alexa Fluor conjugated secondary antibody (Invitrogen). After a further three washes, cells were labelled with DAPI (4,6-diamidino-2-phenylindole; Life Technologies, D1306) for two minutes, washed in PBS, and mounted on glass slides in MOWIOL 4-88 (Calbiochem #475904). For immunolabelling of extracellular matrix, cells were PFA fixed and stained without permeabilization. Collagen and fibronectin fibrillar features were analysed using the matrix analysis Fiji macro TWOMBLI (Wershof et al., 2021).

Widefield images of fixed cells were taken using an Olympus IX70 microscope with 60x 1.42 NA oil-immersion lens, Exfo 120 metal halide illumination with excitation, dichroic and emission filters (Semrock, Rochester, NY), and a Photometrics Coolsnap HQ2 CCD, controlled by Volocity 5.4.1 (Perkin Elmer). Chromatic shifts in images were registration corrected using TetraSpek fluorescent beads (Thermo Fisher). Images were acquired as Δ0.2 µm *z*-stacks.

Confocal microscopy was performed using one of two systems. One, a Leica SP5II AOBS (Acousto-Optical Beam Splitter) confocal laser scanning microscope equipped with 50mW 405nm laser, 150 mW Argon laser, 20mW solid state yellow laser, 20 mW Red He/Ne diode laser, photomultiplier tube detectors and a 63x HCX PL APO CS oil objective. Or two, a Leica SP8 AOBS (Acousto-Optical Beam Splitter) confocal laser scanning microscope equipped with 65 mW Argon laser, 20 mW DPSS yellow laser, 10 mW Red He/Ne and 50 mW 405 nm diode laser, hybrid detectors, and a 63x HC PL APO CS2 oil objective. Images were acquired at 1024 × 1024 x-y resolution, averaging three line scans per channel, using LAS X software. Image stacks were taken with Δz of 0.2 µm.

Golgi area and fragmentation were measured from confocal images using an image J macro with the following procedures: Subtract background (rolling ball size 20), apply median filter (radius determined individual experiments), apply threshold (set for each experiment), make binary, analyse particles (size >0.1).

### Electron microscopy

Cells were grown to confluence in 35mm dishes and then fixed *in situ* in 2.5% glutaraldehyde/0.1 M cacodylate buffer for 20 minutes. Cells were then washed with 0.1 M cacodylate buffer for 10 minutes and fixed in 1% OsO4, 1.5% potassium ferro-cyanide, 0.1 M cacodylate buffer for 60 minutes. Cells were then washed with 0.1 M cacodylate buffer twice, and with water twice for 10 minutes each. To negative stain, cells were incubated with 3% uranyl acetate for 20 minutes and washed again with water. Samples were then dehydrated by successive 10-minute incubations with 70%, 80%, 90%, 96% and 100% EtOH. After a second 100% EtOH incubation, all EtOH was removed and approximately 1ml of a 50:50 mix of propylene oxide and epon was added to the dish. Samples were placed on a rocker for two hours. This was exchanged with 100% epon before placing resin stubs in the dish and baking the samples at 70°C for two days. Stubs were extracted from the dish and trimmed. Ultrathin sections of 70 nm were cut using a diamond knife and a UC6 ultramicrotome (Leica Microsystems) and incubated with 10 nm gold nanoparticles to act as fiducials (Sigma-Aldrich #752584). Sections were imaged using a Tecnai20 LaB6 200k kV twin lens transmission electron microscope (FEI) to collect a tilt series between −70° and +70° using a dedicated Fischione tomography holder and FEI software. Tomograms were reconstructed using IMOD© software. Alignment was computed using fiducial tracking and tomograms generated using 10 iterations with SIRT-like filter reconstruction. Segmentation was performed using Amira 2019.3 software.

### AFM

Cells were grown on coverslips and the cell derived matrix processed as above and then fixed in 4% PFA (Fisher-Scientific P/0840/53) for 10 minutes at room temperature. Coverslips were rinsed with deionised water to remove salt crystals visible in the HS-AFM images and mounted onto SEM stubs. Excess water was blown off with compressed gas and samples were stored at room temperature prior to HS-AFM imaging. Imaging was performed with a HS-AFM (Bristol Nanodynamics) and a MSNL −10 cantilever chip (Bruker). One hundred images in 10 x 10 raster scans were collected in three different areas of each coverslip. Imaging and analysis of the samples was performed blind. Fibrillar features were analysed from the images of one raster scan from each biological replicate (n=3) with the matrix analysis Fiji macro TWOMBLI (Wershof et al., 2021). Identical TWOMBLI parameters were used for all samples. Periodicity was determined by extracting the peak profiles along fibrillar structures using Gwyddion 2.60. The period was calculated using Excel and data displayed/analysed with Graphpad Prism 10.0.2.

### Live imaging of RUSH assays

For live imaging RUSH experiments, cells stably expressing BGN-SBP-mSc were grown to confluence in 35 mm Mattek glass bottom imaging dishes (MatTek P35G-1.5-20C) and transiently transfected with ManII-BFP_IRES_KDEL-streptavidin (McCaughey et al., 2021) 24 hours prior to imaging. To image, growth media was replaced with 1 ml pre-warmed Fluorobrite DMEM (Thermo-Fisher Scientific; A1896701) and dishes were mounted on an Olympus IXplore spinning disk system in a light excluding environmental control chamber (PECON) at 37°C. At T0, 1 ml fluorobrite media containing 80 µM biotin was added to the dish (final concentration 40 µm) and imaging immediately started, taking 120 frames every 10 seconds followed by 4800 frames every 500 ms. Data were acquired using a 60x oil immersion lens with 1.5 numerical aperture, Hamamatsu Fusion BT sCMOS cameras, diode lasers, TruFocus drift compensation, and Olympus CellSens imaging software. The frame at which mScarlet signal could first be seen accumulating adjacent to the BFP signal was used to measure transit to Golgi time. The frame in which a tubular carrier could first be seen emerging from the BFP signal (that then continues to the periphery) was used to calculate Golgi transit time.

### Secretion assays

To perform secretion assays, cells were seeded in 6 well dishes and grown to confluence before aspirating their growth medium and replacing this with 1ml serum-free DMEM F12-HAM. After overnight incubation at 37°C/5% CO2, cells were put on ice, the media was collected into eppendorfs and the cell layer washed with ice cold PBS. Cells were then lysed in RIPA buffer (50 mM Tris-HCl, pH 7.5, 300 mM NaCl, 2% Triton X-100, 1% deoxycholate, 0.1% SDS, 1 mM EDTA) containing protease inhibitors (Millipore #539137) on ice for 15 minutes on a rocker. Media samples were centrifuged at 2000 xg for 2 minutes at 4°C to remove dead cells and the supernatant collected. RIPA lysates were scraped up into eppendorfs and centrifuged for 10 minutes at 13000 xg/4°C and the supernatant collected. Media and lysates were then mixed with 4x Bolt™ LDS sample buffer (Invitrogen™ #B0007), boiled for 10 minutes at 70°C and run on SDS-PAGE gels as below.

For nocodazole secretion assays, cell media was replaced with 1ml serum-free DMEM F12-HAM supplemented with 20 µM nocodazole or an equivalent volume of DMSO as vehicle control for 5 hours. Beyond this point cells were beginning to round and longer collections were not possible. Media and lysates were then collected as normal.

For normal cell lysate collection and western blotting, cells were lysed in RIPA buffer and processed as above. For cell-derived matrix samples, prewarmed sample buffer with reducing agent was added to the dish after cell extraction (above), the dishes were scraped and the buffer collected and boiled for 10 minutes at 70°C.

### Deglycosylation assays

WT and KO cells stably expressing BGN-SBP-mSc were grown to confluence in a 150 mm dish and media exchanged for serum free media 16 hours prior to experiment. Media was then collected into falcon tubes on ice and spun at 1200 rpm for 2 minutes to pellet out cells. The supernatant was collected, supplemented with a protease inhibitor cocktail (Millipore #539137) and BGN-SBP-mSc extracted by RFP trap. To trap, a 60 µl aliquot of Chromotek RFP-trap® agarose beads (Chromotek #rta) was washed 3x in a falcon tube with dilution buffer (10mM Tris/Hcl pH7.4, 50mM NaCl, 0.5mM EDTA), centrifuging for 2 minutes at 2000 rpm at 4°C each time to pellet the beads for buffer exchange. On the final wash, all buffer was removed and the media supernatant added to the beads. Beads and sample were mixed on a rotator at 4°C for 2 hours. Tubes were spun at 2700 xg for 2 minutes at 4°C to pellet the beads and the media removed. Beads were then washed 3x as before in dilution buffer supplemented with protease inhibitors. After the removal of the final wash, the beads were purged.

*For digest with PNGase (kit, NEB P0704S)*, 10x glycoprotein denaturation buffer was diluted to 1x with water and then 80 µl of this added to the bead pellet. Beads and buffer were transferred to an Eppendorf tube and then heated at 100°C for 10 minutes. Samples were then put on ice for 10 seconds before centrifuging them at 2700 xg for 2 minutes. A 60 µl aliquot of the supernatant was transferred to a new tube to act as an undigested control. The remaining 20 µl were transferred to a new tube and contents were digested by adding 4 µl 2x Glycobuffer 2, 4 µl 10% NP40, 10 µl of water, and 2 µl of PNGase F and incubating the mix at 37°C for one hour. Samples were cooled to room temperature, mixed with 4x Bolt™ LDS sample buffer (Invitrogen™ #B0007) and analysed by SDS-PAGE as below.

*For digest with Chondroitinase ABC (Merck Life Science Limited)*, bead pellets were resuspended in 200 µl dilution buffer (50 mM Tris HCl pH8, 60 mM sodium acetate, 0.02% BSA) and heated at 65°C for 30 minutes to heat inactivate any residual proteases. Of this suspension, 100 µl was then taken and 0.3U chondroitinase ABC added (digested sample). The remaining 100 µl was left untreated (undigested). All samples were incubated at 37°C overnight. 4x Bolt™ LDS sample buffer (Invitrogen™ #B0007) was added, and samples boiled at 70°C ready for western blotting as below.

### Western blotting

Samples were run in Bolt 4-12% bis-tris gels with MOPS running buffer for 40 minutes at 130 V. They were then transferred to 0.2 µM nitrocellulose membranes and blocked with 5% milk-TBST. Primary antibody incubations were run overnight at 4°C, then membranes were washed with TBS-0.05% tween (Sigma-Aldrich) and incubated with HRP- (Jackson ImmunoResearch) or fluorophore-conjugated (Invitrogen) secondary antibodies for 2 hours prior to imaging by enhanced chemiluminescence (Promega ECL) and autoradiography films (GE Healthcare) or by fluorescence imaging (LiCor Odyssey) respectively. For collagen blotting, samples were ran on NuPAGE 3-8% tris-acetate gels at 120 V for 1 hour and transferred onto 0.45 µm PVDF membranes overnight at 12 V and 4 °C. Membrane total protein staining was performed with a total protein stain kit (Licor). Quantification of band intensity was performed with image J or Empiria Studio 3.0.

### Matrisome proteomics

Cells were grown in 15 cm dishes for one week in the presence of 50 μg/ml L-ascorbic acid-2- phosphate (Sigma-Aldrich A8960). Cells were then extracted and the cell derived matrix was collected by adding 1 ml PBS to the plate and scraping for 30 seconds. Samples were transferred to tubes and frozen at −20°C prior to processing.

Lysis buffer (10% sodium dodecyl sulfate (Sigma Aldrich) in 50 mM TEAB (Sigma-Aldrich) supplemented with protease and phosphatase inhibitors) was added (1:1 v/v). Samples were sonicated using a Covaris LE220+ sonicator using a 40 W lysis programme with 100 cycles per burst, a duty factor of 40% and a peak incident power of 500 W. Next, 50 µg of protein was reduced and alkylated using DTT (final concentration 5 mM) and IAA (final concentration 15 mM) respectively. Samples were acidified with H_3_PO_4_ (1.2% final concentration), then 7 volumes of Binding buffer (90% methanol, 10% distilled water; 100 mM TEAB) was added and the samples were loaded onto S-Trap columns (ProtiFi) according to the manufacturer’s protocol.

Column-bound proteins were washed in Binding buffer and then digested with 5 µg of 0.8 µg/µL trypsin solution (Promega) diluted in Digestion buffer (50 mM TEAB at pH 8.5). Peptides were eluted from the column in 65 µL Digestion buffer, 65 µl 0.1% formic acid (FA) in distilled water and finally with 30 µL 0.1% FA, 30% acetonitrile (ACN, Sigma). Peptides were desalted using Oligo R3 resin beads (Thermo Scientific), according to the manufacturer’s protocol, in a 96-well, 0.2 µm polyvinylidene fluoride (PVDF) filter plate (Corning). The immobilised peptides were washed twice with 0.1% FA prior to 2 x 50 µl elutions in 0.1% FA with 30% ACN, and lyophilisation using a speed-vac (Heto Cooling System).

Peptides were resuspended in 10 µL 0.1% FA in 5% ACN. Liquid chromatography (LC) separation was performed on a Thermo RSLC system consisting of an NCP3200RS nano pump, WPS3000TPS autosampler and TCC3000RS column oven configured with buffer A as 0.1% FA in water and buffer B as 0.1% DA in ACN. The analytical column (Waters nanoEase M/Z Peptide CSH C18 Column, 130 Å, 1.7 µm, 75 µm x 250 mm) was kept at 35°C and at a flow rate of 300 nL/minute for 8 minutes. The injection valve was set to load before a separation consisting of a 105 minute multistage gradient ranging from 2 to 65% of buffer B. The LC system was coupled to a Thermo Exploris 480 mass spectrometry system via a Thermo Nanospray Flex ion source. The nanospray voltage was set at 1900 V and the ion transfer tube temperature set to 275°C. Data was acquired in a data-dependent manner using a fixed cycle time of 2 seconds, an expected peak width of 15 seconds and a default charge state of +2. Full mass spectra were acquired in positive mode over a scan range of 300 to 1750 Th, with a resolution of 120000, a normalised automatic gain control (AGC) target of 300% and a max fill time of 25 ms for a single microscan. Fragmentation data was obtained from signals with a charge state of +2 or +3 with an intensity over 5000. They were then dynamically excluded from further analysis for a period of 15 seconds after a single acquisition within a 10 ppm window. Fragmentation spectra were acquired with a resolution of 15000, a normalised collision energy of 30%, an AGC target of 300%, a first mass of 110 Th, and a maximum fill time of 25 ms for a single microscan. All data was collected in profile mode.

Mass spectrometry data was analysed using MaxQuant (v1.6.14) using default parameters. All searches included the fixed modification for carbamidomethylation on cysteine residues resulting from IAA treatment. The variable modifications included in the search were oxidised methionine (monoisotopic mass change, +15.955 Da), hydroxyproline (15.995 Da), acetylation of the protein N-terminus (42.011 Da) and phosphorylation of threonine, serine and tyrosine (79.966 Da). A maximum of 2 missed cleavages per peptide was allowed. Peptides were searched against the UniProt database with a maximum false discovery rate of 1%. Proteins were required to have a minimum false discovery rate (FDR) of 1% and at least 2 unique peptides in order to be accepted, known contaminants were also removed. Missing values were assumed to be due to low abundance. Abundances were compared using MsqRob where label free quantification data was normalised using median peptide intensity (Goeminne et al., 2020). MsqRob was used using R (version 4.1.2). The remaining statistics were done using Prism (GraphPad Software, version 9.1.2).

### Secretome proteomics

Cells were seeded into 6 well dishes and grown to confluence. Cells were rinsed once with serum free medium, taking care to aspirate as much media as possible each time. Exactly 1 ml serum free F12 HAM was then added to cells overnight. After 24 hours, cells were put on ice and 900 µl medium was removed from well and added to eppendorf on ice. Media were spun at 2000 rpm for 5 minutes and then 800 µl removed and put in a fresh tube on ice. To test protein concentration, 25 µl medium was used in a BSA assay prior to snap freezing and storage at −80 °C.

For TMT Labelling and High pH reversed-phase chromatography, an equal volume (95 µl) of each sample was digested with trypsin (1.25 µg trypsin; 37 °C, overnight), labelled with Tandem Mass Tag (TMTpro) sixteen plex reagents according to the manufacturer’s protocol (Thermo Fisher Scientific, Loughborough, LE11 5RG, UK) and the labelled samples pooled. The pooled samples were desalted using a SepPak cartridge according to the manufacturer’s instructions (Waters, Milford, Massachusetts, USA). Eluate from the SepPak cartridge was evaporated to dryness and resuspended in buffer A (20 mM ammonium hydroxide, pH 10) prior to fractionation by high pH reversed-phase chromatography using an Ultimate 3000 liquid chromatography system (Thermo Fisher Scientific). In brief, the sample was loaded onto an XBridge BEH C18 Column (130Å, 3.5 µm, 2.1 mm X 150 mm, Waters, UK) in buffer A and peptides eluted with an increasing gradient of buffer B (20 mM Ammonium Hydroxide in acetonitrile, pH 10) from 0-95% over 60 minutes. The resulting fractions (concatenated into 15 in total) were evaporated to dryness and resuspended in 1% formic acid prior to analysis by nano-LC MSMS using an Orbitrap Fusion Lumos mass spectrometer (Thermo Scientific).

High pH RP fractions were further fractionated using an Ultimate 3000 nano-LC system in line with an Orbitrap Fusion Lumos mass spectrometer (Thermo Scientific). In brief, peptides in 1% (vol/vol) formic acid were injected onto an Acclaim PepMap C18 nano-trap column (Thermo Scientific). After washing with 0.5% (vol/vol) acetonitrile 0.1% (vol/vol) formic acid peptides were resolved on a 250 mm × 75 μm Acclaim PepMap C18 reverse phase analytical column (Thermo Scientific) over a 150 min organic gradient, using 7 gradient segments (1-6% solvent B over 1min., 6-15% B over 58min., 15-32% B over 58min., 32-40% B over 5min., 40-90% B over 1min., held at 90% B for 6min and then reduced to 1% B over 1min.) with a flow rate of 300 nl min^−1^. Solvent A was 0.1% formic acid (FA) and Solvent B was aqueous 80% acetonitrile in 0.1% formic acid. Peptides were ionized by nano-electrospray ionization at 2.0 kV using a stainless-steel emitter with an internal diameter of 30 μm (Thermo Scientific) and a capillary temperature of 300 °C.

All spectra were acquired using an Orbitrap Fusion Lumos mass spectrometer controlled by Xcalibur 3.0 software (Thermo Scientific) and operated in data-dependent acquisition mode using an SPS-MS3 workflow. FTMS1 spectra were collected at a resolution of 120 000, with an automatic gain control (AGC) target of 200 000 and a max injection time of 50 ms. Precursors were filtered with an intensity threshold of 5000, according to charge state (to include charge states 2-7) and with monoisotopic peak determination set to Peptide. Previously interrogated precursors were excluded using a dynamic window (60s +/-10ppm). The MS2 precursors were isolated with a quadrupole isolation window of 0.7m/z. ITMS2 spectra were collected with an AGC target of 10 000, max injection time of 70ms and CID collision energy of 35%.

For FTMS3 analysis, the Orbitrap was operated at 50 000 resolution with an AGC target of 50 000 and a max injection time of 105 ms. Precursors were fragmented by high energy collision dissociation (HCD) at a normalised collision energy of 60% to ensure maximal TMT reporter ion yield. Synchronous Precursor Selection (SPS) was enabled to include up to 10 MS2 fragment ions in the FTMS3 scan.

The raw data files were processed and quantified using Proteome Discoverer software v2.1 (Thermo Scientific) and searched against the UniProt Human database (downloaded January 2023: 81579 entries) and the Uniprot Bos taurus database (downloaded February 2023: 37505 entries) using the SEQUEST HT algorithm. Peptide precursor mass tolerance was set at 10ppm, and MS/MS tolerance was set at 0.6Da. Search criteria included oxidation of methionine (+15.995Da), acetylation of the protein N-terminus (+42.011Da) and Methionine loss plus acetylation of the protein N-terminus (- 89.03Da) as variable modifications and carbamidomethylation of cysteine (+57.021Da) and the addition of the TMTpro mass tag (+304.207) to peptide N-termini and lysine as fixed modifications. Searches were performed with full tryptic digestion and a maximum of 2 missed cleavages were allowed. The reverse database search option was enabled and all data was filtered to satisfy false discovery rate (FDR) of 5%.

As samples are from the media fraction of cultures, equal volumes of sample were analysed rather than equal quantities of protein, and raw abundances were used without normalisation. Statistical analysis was performed in R version 4.3.0. PCAs were calculated using the FactoMineR package, PCAs, Sum abundance, and volcano plots were plotted in ggplot2. A linear mixed effect model was fitted for each protein using the package lme4, with replicate number included as a random effect, and both clones included in a single variable for the KO condition to increase statistical power. P-values were then FDR adjusted using the Benjamini Hochberg method to account for multiple testing. Note, this was used as a gold-standard metric for individual proteins rather than a threshold for statistical analysis, as it is understood to be an overly stringent method for most proteomics experiments (Pascovici et al., 2016). Downstream analysis was performed using QIAGEN IPA (QIAGEN Inc., https://digitalinsights.qiagen.com/IPA), using p<0.05 as a cut-off and the unfiltered user data as the reference set (Kramer et al., 2014). GO term functional enrichment analysis was performed using the Database for Annotation, Visualization, and Integrated Discovery (DAVID; http://www.david.niaid.nih.gov).

### qPCR

Cells were grown to confluence in 35 mm dishes and then RNA was extracted using RNeasy® plus mini kit (QIAGEN #74136). Reverse transcription was performed using an Invitrogen Superscript III kit (Invitrogen™ #11752-050) following the manufacturer’s instructions to generate cDNA. cDNA was diluted to 15 ng/µl and primers to 2 µM stocks. Reactions were set up using 3.75 ng/µl DNA, 500 nM forward and reverse primers, and 2x DyNAmo Flash Sybr green master mix (ThermoFisher Scientific #F415) in MicroAmp® optical 96-well reaction plates (Applied biosystems #N8010560). RT-PCR was then performed on a QuantStudio™ 3 real time PCR system using cycle: 1) 7 minutes @ 95°C, 2) 15 seconds @ 95°C, 3) 45 seconds @ 60°C, 4) 40 cycles of steps 2-3. RT-PCR was followed by melting curve analysis at 60-98°C for quality control. PCR products were tested on a gel for size during primer validation. RT-PCR results were analysed with the QuantStudio™ real time PCR software.

**Primer sequences.**
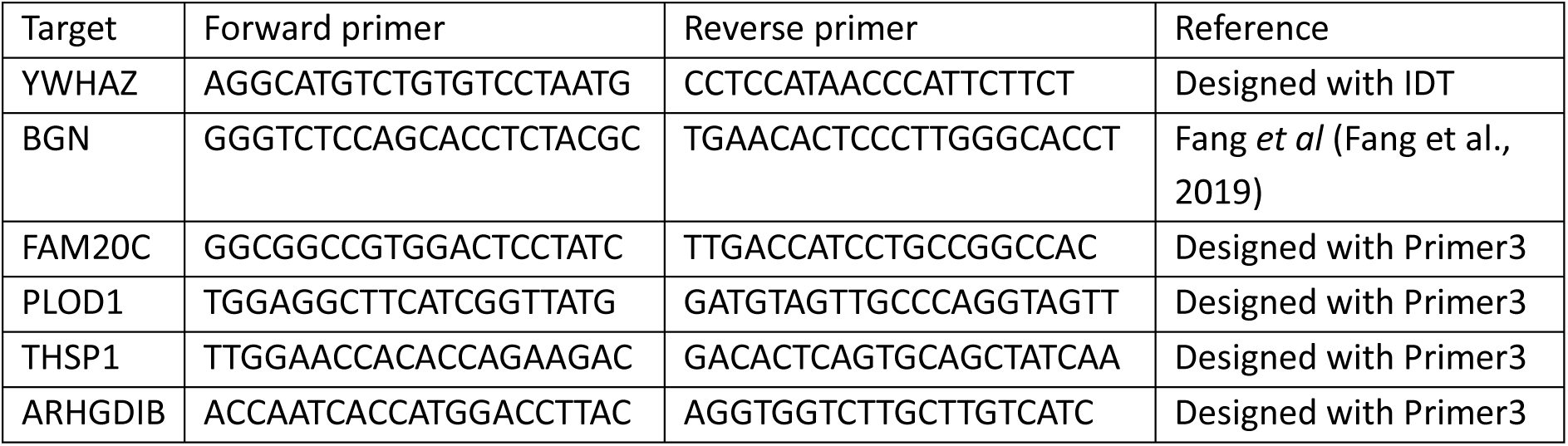

### Statistical analysis

All statistical analyses were performed with graphpad prism. Data were subjected to normality testing using the Shapiro-Wilk test. Data found to be normally distributed were analysed by t-test or one-way ANOVA and the mean and standard deviation shown. Non-normally distributed data were analysed with a Mann-Whiney test, or to compare multiple conditions Kruskall-Wallis test with Dunn’s multiple comparisons, and the median and interquartile range presented. Statistical information is provided in each figure legend.

## Acknowledgements

We would like to thank Andrew Herman and Poppy Miller of the UoB FACS facility with their help sorting CRISPR transfected cells, the UoB Wolfson facility, especially Judith Mantell and Dominic Alibhai, for training advice and facilities, David Knight and Stacey Harwood at the University of Manchester BioMS facility for running the matrisome samples, Prof Oliver Jenson and Dr Christopher Revell for conceptual discussions, Dr Anne George and Dr Harry Young for running pilot experiments, and Prof Stuart Haslam for advice. This work was funded by the UK Research and Innovation– Biotechnology and Biological Sciences Research Council BB/T001984/1.

## CRediT Author contributions

Conceptualisation: D.J.S, N.L.S, M.L.; Methodology: N.L.S, G.T, A.H, K.H, P.A.L; Validation: N.L.S, G.T.; Formal analysis: N.L.S, P.A.L, A.H, G.T.; Investigation: N.L.S, G.T, A.H, E.P.S, K.H.; Writing draft: N.L.S, G.T, A.H, P.A.L; Writing review and editing: N.L.S, M.L; Visualisation: N.L.S., G.T, A.H, P.A.L; Supervision: N.L.S, D.J.S, M.L., J.S; Project administration: N.L.S, D.J.S; Funding acquisition: D.J.S, M.L, N.L.S, J.S.

## References

Adams, J.C. 2023. Passing the post: roles of posttranslational modifications in the form and function of extracellular matrix. Am J Physiol Cell Physiol. 324:C1179–C1197.

Adusumalli, R., H.C. Asheim, V. Lupashin, J.B. Blackburn, and K. Prydz. 2021. Proteoglycan synthesis in conserved oligomeric Golgi subunit deficient HEK293T cells is affected differently, depending on the lacking subunit. Traffic. 22:230–239.

Ahat, E., Y. Song, K. Xia, W. Reid, J. Li, S. Bui, F. Zhang, R.J. Linhardt, and Y. Wang. 2022. GRASP depletion-mediated Golgi fragmentation impairs glycosaminoglycan synthesis, sulfation, and secretion. Cell Mol Life Sci. 79:199.

Appenzeller-Herzog, C., and H.P. Hauri. 2006. The ER-Golgi intermediate compartment (ERGIC): in search of its identity and function. J Cell Sci. 119:2173–2183.

Arab, M., T. Chen, and M. Lowe. 2024. Mechanisms governing vesicle traffic at the Golgi apparatus. Curr Opin Cell Biol. 88:102365.

Arpino, V., M. Brock, and S.E. Gill. 2015. The role of TIMPs in regulation of extracellular matrix proteolysis. Matrix Biol. 44–46:247-254.

Banu, Y., M. Matsuda, M. Yoshihara, M. Kondo, S. Sutou, and S. Matsukuma. 2002. Golgi matrix protein gene, Golga3/Mea2, rearranged and re-expressed in pachytene spermatocytes restores spermatogenesis in the mouse. Mol Reprod Dev. 61:288–301.

Barr, F.A., and B. Short. 2003. Golgins in the structure and dynamics of the Golgi apparatus. Curr Opin Cell Biol. 15:405–413.

Bentson, L.F., V.A. Agbor, L.N. Agbor, A.C. Lopez, L.E. Nfonsam, S.S. Bornstein, M.A. Handel, and C.C. Linder. 2013. New point mutation in Golga3 causes multiple defects in spermatogenesis. Andrology. 1:440–450.

Bergen, D.J.M., N.L. Stevenson, R.E.H. Skinner, D.J. Stephens, and C.L. Hammond. 2017. The Golgi matrix protein giantin is required for normal cilia function in zebrafish. Biol Open. 6:1180–1189.

Bird, I.M., S.H. Kim, D.K. Schweppe, J. Caetano-Lopes, A.G. Robling, J.F. Charles, S.P. Gygi, M.L. Warman, and P.J. Smits. 2018. The skeletal phenotype of achondrogenesis type 1A is caused exclusively by cartilage defects. Development. 145.

Boncompain, G., L. Fourriere, N. Gareil, and F. Perez. 2021. Retention Using Selective Hooks-Synchronized Secretion to Measure Local Exocytosis. Methods Mol Biol. 2233:253–264.

Chang, W.L., C.W. Chang, Y.Y. Chang, H.H. Sung, M.D. Lin, S.C. Chang, C.H. Chen, C.W. Huang, K.S. Tung, and T.B. Chou. 2013. The Drosophila GOLPH3 homolog regulates the biosynthesis of heparan sulfate proteoglycans by modulating the retrograde trafficking of exostosins. Development. 140:2798–2807.

Costantini, A., H. Valta, A.M. Suomi, O. Makitie, and F. Taylan. 2021. Oligogenic Inheritance of Monoallelic TRIP11, FKBP10, NEK1, TBX5, and NBAS Variants Leading to a Phenotype Similar to Odontochondrodysplasia. Front Genet. 12:680838.

Couchman, J.R., and C.A. Pataki. 2012. An introduction to proteoglycans and their localization. J Histochem Cytochem. 60:885–897.

Del Pino, M., M.J. Sanchez-Soler, M. Parron-Pajares, M. Aza-Carmona, K.E. Heath, and V. Fano. 2021. Description of four patients with TRIP11 variants expand the clinical spectrum of odontochondroplasia (ODCD) and demonstrate the existence of common variants. Eur J Med Genet. 64:104198.

Follit, J.A., J.T. San Agustin, F. Xu, J.A. Jonassen, R. Samtani, C.W. Lo, and G.J. Pazour. 2008. The Golgin GMAP210/TRIP11 anchors IFT20 to the Golgi complex. PLoS Genet. 4:e1000315.

Gandhi, N.S., and R.L. Mancera. 2008. The structure of glycosaminoglycans and their interactions with proteins. Chem Biol Drug Des. 72:455–482.

Gillingham, A.K., and S. Munro. 2016. Finding the Golgi: Golgin Coiled-Coil Proteins Show the Way. Trends Cell Biol. 26:399–408.

Guinec, N., V. Dalet-Fumeron, and M. Pagano. 1993. “In vitro” study of basement membrane degradation by the cysteine proteinases, cathepsins B, B-like and L. Digestion of collagen IV, laminin, fibronectin, and release of gelatinase activities from basement membrane fibronectin. Biol Chem Hoppe Seyler. 374:1135–1146.

Harada, A., M. Kunii, K. Kurokawa, T. Sumi, S. Kanda, Y. Zhang, S. Nadanaka, K.M. Hirosawa, K. Tokunaga, T. Tojima, M. Taniguchi, K. Moriwaki, S.I. Yoshimura, M. Yamamoto-Hino, S. Goto, T. Katagiri, S. Kume, M. Hayashi-Nishino, M. Nakano, E. Miyoshi, K.G.N. Suzuki, H. Kitagawa, and A. Nakano. 2024. Dynamic movement of the Golgi unit and its glycosylation enzyme zones. Nat Commun. 15:4514.

Hellicar, J., N.L. Stevenson, D.J. Stephens, and M. Lowe. 2022. Supply chain logistics - the role of the Golgi complex in extracellular matrix production and maintenance. J Cell Sci. 135.

Hicks, S.W., T.A. Horn, J.M. McCaffery, D.M. Zuckerman, and C.E. Machamer. 2006. Golgin-160 promotes cell surface expression of the beta-1 adrenergic receptor. Traffic. 7:1666–1677.

Hicks, S.W., and C.E. Machamer. 2005. Isoform-specific interaction of golgin-160 with the Golgi-associated protein PIST. J Biol Chem. 280:28944–28951.

Jangra, J., N.G. Bajad, R. Singh, A. Kumar, and S.K. Singh. 2024. Identification of novel potential cathepsin-B inhibitors through pharmacophore-based virtual screening, molecular docking, and dynamics simulation studies for the treatment of Alzheimer’s disease. Mol Divers.

Karamanos, N.K., A.D. Theocharis, Z. Piperigkou, D. Manou, A. Passi, S.S. Skandalis, D.H. Vynios, V. Orian-Rousseau, S. Ricard-Blum, C.E.H. Schmelzer, L. Duca, M. Durbeej, N.A. Afratis, L. Troeberg, M. Franchi, V. Masola, and M. Onisto. 2021. A guide to the composition and functions of the extracellular matrix. FEBS J. 288:6850–6912.

Katayama, K., M. Kuriki, T. Kamiya, Y. Tochigi, and H. Suzuki. 2018. Giantin is required for coordinated production of aggrecan, link protein and type XI collagen during chondrogenesis. Biochem Biophys Res Commun. 499:459–465.

Katayama, K., T. Sasaki, S. Goto, K. Ogasawara, H. Maru, K. Suzuki, and H. Suzuki. 2011. Insertional mutation in the Golgb1 gene is associated with osteochondrodysplasia and systemic edema in the OCD rat. Bone. 49:1027–1036.

Kikukawa, K., T. Kamei, and K. Suzuki. 1991a. Chromatographic analysis of glycosaminoglycans in epiphyseal cartilage of congenital osteochondrodysplasia (ocd/ocd) rat. J Vet Med Sci. 53:1091–1092.

Kikukawa, K., T. Kamei, and K. Suzuki. 1991b. A histological and histochemical study on glycosaminoglycans in epiphysial cartilage of osteochondrodysplasia rat (OCD/OCD). Connect Tissue Res. 25:301–309.

Kikukawa, K., T. Kamei, K. Suzuki, and K. Maita. 1990. Electron microscopic observations and electrophoresis of the glycosaminoglycans in the epiphyseal cartilage of the congenital osteochondrodysplasia rat (ocd/ocd). Matrix. 10:378–387.

Kikukawa, K., and K. Suzuki. 1992. Histochemical and immunohistochemical distribution of glycosaminoglycans, type II collagen, and fibronectin in developing fetal cartilage of congenital osteochondrodysplasia rat (ocd/ocd). Teratology. 46:509–523.

Kramer, A., J. Green, J. Pollard, Jr., and S. Tugendreich. 2014. Causal analysis approaches in Ingenuity Pathway Analysis. Bioinformatics. 30:523–530.

Lan, Y., N. Zhang, H. Liu, J. Xu, and R. Jiang. 2016. Golgb1 regulates protein glycosylation and is crucial for mammalian palate development. Development. 143:2344–2355.

Lazaro-Dieguez, F., N. Jimenez, H. Barth, A.J. Koster, J. Renau-Piqueras, J.L. Llopis, K.N. Burger, and G. Egea. 2006. Actin filaments are involved in the maintenance of Golgi cisternae morphology and intra-Golgi pH. Cell Motil Cytoskeleton. 63:778–791.

Lee, M.H., and G. Murphy. 2004. Matrix metalloproteinases at a glance. J Cell Sci. 117:4015–4016.

Lohmander, S., K. Madsen, and A. Hinek. 1979. Secretion of proteoglycans by chondrocytes. Influence of colchicine, cytochalasin B, and beta-D-xyloside. Arch Biochem Biophys. 192:148–157.

Lowe, M. 2011. Structural organization of the Golgi apparatus. Curr Opin Cell Biol. 23:85–93.

Lowe, M. 2019. The Physiological Functions of the Golgin Vesicle Tethering Proteins. Front Cell Dev Biol. 7:94.

Matsukuma, S., M. Kondo, M. Yoshihara, M. Matsuda, T. Utakoji, and S. Sutou. 1999. Mea2/Golga3 gene is disrupted in a line of transgenic mice with a reciprocal translocation between Chromosomes 5 and 19 and is responsible for a defective spermatogenesis in homozygotes. Mamm Genome. 10:1–5.

McCaughey, J., N.L. Stevenson, S. Cross, and D.J. Stephens. 2019. ER-to-Golgi trafficking of procollagen in the absence of large carriers. J Cell Biol. 218:929–948.

McCaughey, J., N.L. Stevenson, J.M. Mantell, C.R. Neal, A. Paterson, K. Heesom, and D.J. Stephens. 2021. A general role for TANGO1, encoded by MIA3, in secretory pathway organization and function. J Cell Sci. 134.

Medina, C.T.N., R. Sandoval, G. Oliveira, K. da Costa Silveira, D.P. Cavalcanti, and R. Pogue. 2020. Pathogenic variants in the TRIP11 gene cause a skeletal dysplasia spectrum from odontochondrodysplasia to achondrogenesis 1A. Am J Med Genet A. 182:681–688.

Pascovici, D., D.C. Handler, J.X. Wu, and P.A. Haynes. 2016. Multiple testing corrections in quantitative proteomics: A useful but blunt tool. Proteomics. 16:2448–2453.

Podgorski, I., B.E. Linebaugh, J.E. Koblinski, D.L. Rudy, M.K. Herroon, M.B. Olive, and B.F. Sloane. 2009. Bone marrow-derived cathepsin K cleaves SPARC in bone metastasis. Am J Pathol. 175:1255–1269.

Potelle, S., A. Klein, and F. Foulquier. 2015. Golgi post-translational modifications and associated diseases. J Inherit Metab Dis. 38:741–751.

Qian, Y., G. Hu, M. Chen, B. Liu, K. Yan, C. Zhou, Y. Yu, and M. Dong. 2021. Novel deep intronic and frameshift mutations causing a TRIP11-related disorder. Am J Med Genet A. 185:2482–2487.

Rabouille, C., N. Hui, F. Hunte, R. Kieckbusch, E.G. Berger, G. Warren, and T. Nilsson. 1995. Mapping the distribution of Golgi enzymes involved in the construction of complex oligosaccharides. J Cell Sci. 108 (Pt 4):1617–1627.

Ricard-Blum, S., R.R. Vives, L. Schaefer, M. Gotte, R. Merline, A. Passi, P. Heldin, A. Magalhaes, C.A. Reis, S.S. Skandalis, N.K. Karamanos, S. Perez, and D. Nikitovic. 2024. A biological guide to glycosaminoglycans: current perspectives and pending questions. FEBS J. 291:3331–3366.

Sato, K., P. Roboti, A.A. Mironov, and M. Lowe. 2015. Coupling of vesicle tethering and Rab binding is required for in vivo functionality of the golgin GMAP-210. Mol Biol Cell. 26:537–553.

Shao, X., C.D. Gomez, N. Kapoor, J.M. Considine, C. Grams, Y.T. Gao, and A. Naba. 2023. MatrisomeDB 2.0: 2023 updates to the ECM-protein knowledge database. Nucleic Acids Res. 51:D1519–D1530.

Smits, P., A.D. Bolton, V. Funari, M. Hong, E.D. Boyden, L. Lu, D.K. Manning, N.D. Dwyer, J.L. Moran, M. Prysak, B. Merriman, S.F. Nelson, L. Bonafe, A. Superti-Furga, S. Ikegawa, D. Krakow, D.H. Cohn, T. Kirchhausen, M.L. Warman, and D.R. Beier. 2010. Lethal skeletal dysplasia in mice and humans lacking the golgin GMAP-210. N Engl J Med. 362:206–216.

Stevenson, N.L., D.J.M. Bergen, Y. Lu, M.E. Prada-Sanchez, K.E. Kadler, C.L. Hammond, and D.J. Stephens. 2021. Giantin is required for intracellular N-terminal processing of type I procollagen. J Cell Biol. 220.

Stevenson, N.L., D.J.M. Bergen, R.E.H. Skinner, E. Kague, E. Martin-Silverstone, K.A. Robson Brown, C.L. Hammond, and D.J. Stephens. 2017. Giantin-knockout models reveal a feedback loop between Golgi function and glycosyltransferase expression. J Cell Sci. 130:4132–4143.

Suzuki, K., K. Kikukawa, Y. Hakamata, T. Kamei, and T. Imamichi. 1988. Congenital osteochondrodysplasia with systemic subcutaneous edema (ocd/ocd): a new lethal autosomal recessive mutant of the rat. J Hered. 79:48–50.

Tie, H.C., D. Mahajan, and L. Lu. 2022. Visualizing intra-Golgi localization and transport by side-averaging Golgi ministacks. J Cell Biol. 221.

Upadhyai, P., P. Radhakrishnan, V.S. Guleria, N. Kausthubham, S.S. Nayak, A. Superti-Furga, and K.M. Girisha. 2021. Biallelic deep intronic variant c.5457+81T>A in TRIP11 causes loss of function and results in achondrogenesis 1A. Hum Mutat. 42:1005–1014.

Vidak, E., U. Javorsek, M. Vizovisek, and B. Turk. 2019. Cysteine Cathepsins and their Extracellular Roles: Shaping the Microenvironment. Cells. 8.

Vizovisek, M., M. Fonovic, and B. Turk. 2019. Cysteine cathepsins in extracellular matrix remodeling: Extracellular matrix degradation and beyond. Matrix Biol. 75–76:141-159.

Waldo, G.S., B.M. Standish, J. Berendzen, and T.C. Terwilliger. 1999. Rapid protein-folding assay using green fluorescent protein. Nat Biotechnol. 17:691–695.

Wehrle, A., T.M. Witkos, S. Unger, J. Schneider, J.A. Follit, J. Hermann, T. Welting, V. Fano, M. Hietala, N. Vatanavicharn, K. Schoner, J. Spranger, M. Schmidts, B. Zabel, G.J. Pazour, A. Bloch-Zupan, G. Nishimura, A. Superti-Furga, M. Lowe, and E. Lausch. 2019. Hypomorphic mutations of TRIP11 cause odontochondrodysplasia. JCI Insight. 4.

Wershof, E., D. Park, D.J. Barry, R.P. Jenkins, A. Rullan, A. Wilkins, K. Schlegelmilch, I. Roxanis, K.I. Anderson, P.A. Bates, and E. Sahai. 2021. A FIJI macro for quantifying pattern in extracellular matrix. Life Sci Alliance. 4.

Williams, D., S.W. Hicks, C.E. Machamer, and J.E. Pessin. 2006. Golgin-160 is required for the Golgi membrane sorting of the insulin-responsive glucose transporter GLUT4 in adipocytes. Mol Biol Cell. 17:5346–5355.

Witkos, T.M., and M. Lowe. 2015. The Golgin Family of Coiled-Coil Tethering Proteins. Front Cell Dev Biol. 3:86.

Wong, M., A.K. Gillingham, and S. Munro. 2017. The golgin coiled-coil proteins capture different types of transport carriers via distinct N-terminal motifs. BMC Biol. 15:3.

Wong, M., and S. Munro. 2014. Membrane trafficking. The specificity of vesicle traffic to the Golgi is encoded in the golgin coiled-coil proteins. Science. 346:1256898.

Yadav, S., M.A. Puthenveedu, and A.D. Linstedt. 2012. Golgin160 recruits the dynein motor to position the Golgi apparatus. Dev Cell. 23:153–165.

Yamaguchi, H., M.D. Meyer, L. He, and Y. Komatsu. 2022. Disruption of Trip11 in cranial neural crest cells is associated with increased ER and Golgi stress contributing to skull defects in mice. Dev Dyn. 251:1209–1222.

Yamaguchi, H., M.D. Meyer, L. He, L. Senavirathna, S. Pan, and Y. Komatsu. 2021. The molecular complex of ciliary and golgin protein is crucial for skull development. Development. 148.

Yeter, B., A. Dilruba Aslanger, G. Yesil, and N.H. Elcioglu. 2022. A Novel Mutation in the TRIP11 Gene: Diagnostic Approach from Relatively Common Skeletal Dysplasias to an Extremely Rare Odontochondrodysplasia. J Clin Res Pediatr Endocrinol. 14:475–480.

Zhang, X., and Y. Wang. 2016. Glycosylation Quality Control by the Golgi Structure. J Mol Biol. 428:3183–3193.

